# Genetic identification of novel medullary neurons underlying congenital central hypoventilation syndrome

**DOI:** 10.1101/2023.05.29.542652

**Authors:** Ke Cui, Yiling Xia, Abhisarika Patnaik, Elijah D. Lowenstein, Eser Göksu Isik, Adrian L. Knorz, Laura Airaghi, Michela Crotti, Michèle Studer, Filippo M. Rijli, Hans G. Nothwang, Luis R. Hernandez-Miranda

**Author notes:** Correspondence should be addressed to: Luis Rodrigo Hernandez-Miranda, Charitéplatz 1 (intern Virchowweg 6), 10117, Berlin, Germany. Tel.: +49 30,450528403. **Co-First authors**. Ke Cui and Yiling Xia shared similar responsibilities and carried out equal amounts of work towards this project. The co-first author order was determined alphabetically.

## Abstract

Congenital Central Hypoventilation Syndrome (CCHS) is a rare, but life-threatening, respiratory disorder that is classically diagnosed in children. This disease is characterized by pronounced alveolar hypoventilation and diminished chemoreflexes, particularly to abnormally high levels of arterial pCO_2_. Mutations in the transcription factors *PHOX2B* and *LBX1* have been identified in CCHS patients, but the dysfunctional circuit responsible for this disease remains unknown. Here, we show that distinct sets of medullary neurons co-expressing both transcription factors (dB2 neurons) account for specific respiratory functions and phenotypes seen in CCHS. By combining murine intersectional chemogenetics, intersectional labeling, and the selective targeting of the CCHS disease-causing *Lbx1^FS^* mutation to specific subgroups of dB2 neurons, we uncovered novel sets of these cells key for i) respiratory tidal volumes and the hypercarbic reflex, ii) neonatal respiratory stability and iii) neonatal survival. These data provide functional evidence for the essential role of dB2 neurons in neonatal respiratory physiology and will be instrumental for the development of therapeutic strategies for the management of CCHS. In summary, our work uncovers new neural components of the central circuit regulating breathing and establishes dB2 neuron dysfunction to be causative of CCHS.

## Introduction

Genetic and environmental factors contribute to the inception of respiratory disorders, but the affected neural circuits are largely unknown. CCHS (OMIM 209880) is a life-threatening disorder with a severe presentation of respiratory and autonomous nervous system dysregulation (1–4). This condition is classically diagnosed in infants and is characterized by primary alveolar hypoventilation while awake and central apnea during sleep (5, 6). Cessation of breathing is less typical than hypoventilation in CCHS patients. However, severely affected children suffer from spontaneous respiratory arrest regardless of their arousal state (7, 8). A conspicuous feature of this disorder is the inability to adjust breathing in response to changes in tissue gas levels, particularly to abnormally high levels of arterial pCO_2_ (9, 10).

CCHS is unique in the sense that clear genetic disturbances have been identified to be causative of this disorder (8, 11–15). Most patients present with dominant *de novo* mutations in *PHOX2B* (8, 13, 16, 17), a gene that encodes for a homeodomain transcription factor that is essential for the development and function of central and peripheral visceral neurons (18–21). Unsurprisingly, patients with *PHOX2B* mutations also have variable manifestations of autonomic nervous system dysregulation, including Hirschsprung’s disease (a rare disorder that produces aganglionosis of the distal hindgut) or neural-crest tumors (4, 5, 22–24). Mouse models carrying human *PHOX2B* mutations die during embryonic life or soon after birth (7, 25–28). To date, the dysfunctional neural circuit responsible for CCHS has not been elucidated. Previously, we identified a recessive frameshift mutation (termed *LBX1^FS^*) in the homeodomain transcription factor *LBX1* to be causative of a severe manifestation of CCHS (12). Unlike PHOX2B that has multiple functions in visceral neurons across the central and peripheral nervous system, the function of LBX1 is limited to specific neuron types in the medulla and spinal cord (29–35). Interestingly, homozygous *Lbx1^FS^* mutant mice recapitulate the severe respiratory phenotypes observed in CCHS and seem to display a unique deficit in the development of at least two medullary neuron groups that co-express both Lbx1 and Phox2b (12).

In mice, *Lbx1* is essential for the specification of four distinct medullary neuron types known as dB1, dB2, dB3 and dB4 (30, 31, 33–35). Notably, dB2 neurons are the only neuron type in the entire nervous system that co-expresses Lbx1 and Phox2b during development (12, 30, 31, 35, 36). These neurons originate from a discrete pool of Phox2b+ progenitor cells called the dB2 progenitor domain, which transiently resides between rhombomeres 2 and 6 (reviewed in 31, 35). As these progenitors become postmitotic, they switch on the expression of *Lbx1* and predictably migrate to distinct locations in the brainstem. Even though most neurons emanating from the dB2 progenitor domain have not been systematically characterized, the conditional restriction of the *Lbx1^FS^* mutation to Phox2b-expressing cells (in *Phox2b^Cre/+^;Lbx1^FS/lox^* mice) recapitulates the severe respiratory phenotype seen in homozygous *Lbx1^FS^* animals (12). This includes pronounced gasping behavior and mortality immediately after birth, although it should be noted that a few mutant mice can survive for up to two hours while displaying robust hypoventilation and marked apneic behavior. This suggests that deficits in dB2 neuron function might be causative of CCHS. However, the premature death of homozygous and conditional *Lbx1^FS^* mice precluded the definitive association of dB2 neuron dysfunction to the disease. Similarly, it is presently unknown whether all or specific subgroups of dB2 neurons (i.e. dB2 neurons originating from distinct rhombomeres) participate in respiration.

In this study, we set out to investigate the potential role of dB2 neurons in respiration and CCHS. Using murine intersectional chemogenetics, we show for the first time that dB2 neurons are critical for the generation of adequate respiratory tidal volumes. Furthermore, our experiments illustrate that dB2 neurons have a prominent role in respiratory frequency patterns, respiratory stability and the hypercarbic reflex in neonates. By restricting the CCHS disease-causing *Lbx1^FS^* variant to specific rhombomeres, we show that a single subgroup of dB2 neurons (from rhombomere 5) is crucial for respiratory frequency and the response to hypercarbia in neonates, while distinct subgroups of dB2 neurons (generated in rhombomere 6) are key for: i) respiratory tidal volumes, ii) neonatal respiratory stability, and iii) neonatal survival. Our work thus identifies new neural components of the central respiratory circuit, whose dysfunction causes all respiratory phenotypes seen in CCHS patients.

## Results

### Silencing of dB2 neurons causes hypoventilation

To directly determine whether dB2 neurons play a function in breathing, we used a murine intersectional chemogenetic strategy to transiently and reversibly silence their neural activity at different postnatal times. To do so, we used the *RC::FPDi* dual-recombinase allele that expresses an HA-tagged hM4Di, a mutant G protein-coupled DREADD receptor, upon the excision of two stop cassettes flanked by *FRT* and *LoxP* sites. These stop cassettes were excised by Cre and FlpO recombinases driven by *Lbx1* (*Lbx1^Cre^*) and *Phox2b* (*Phox2b^FlpO^*), respectively (Figures 1A and 1B left).

**Figure 1.**
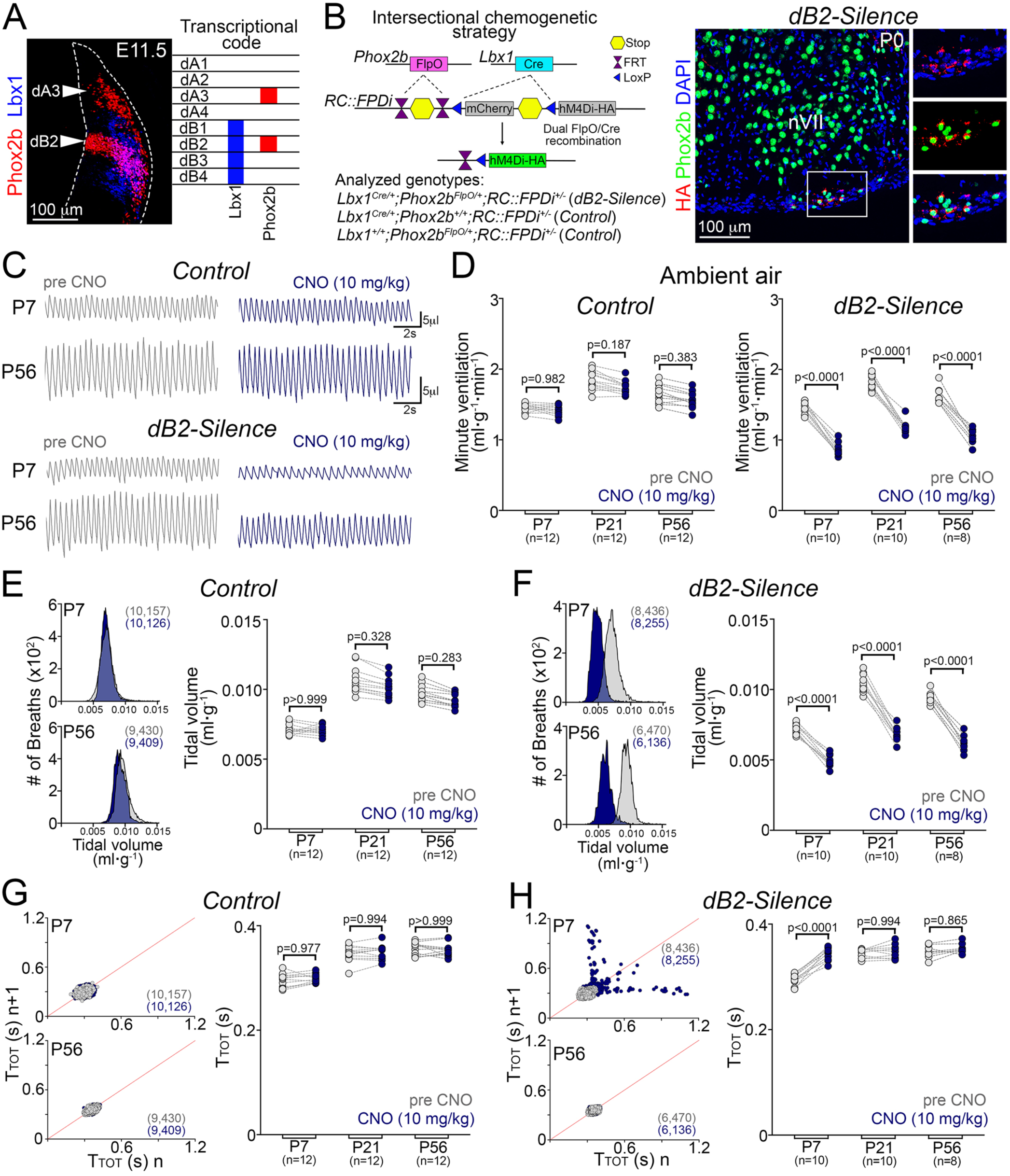
Silencing of dB2 neurons causes hypoventilation. (**A**) Left, a transverse brainstem section stained with Phox2b and Lbx1 antibodies at embryonic (E) day 11.5. Right, scheme illustrating genes expressed by brainstem neurons at E11.5. Note that Lbx1 and Phox2b are co-expressed only in dB2 neurons. (**B**) Left, intersectional genetic strategy to express the hM4Di-HA DREADD receptor in dB2 neurons. Analyzed genotypes are indicated. Right, a transverse brainstem section stained with Phox2b and HA antibodies at birth (P0). The facial motor nucleus (nVII) is indicated. DAPI was used to counterstain. Note that only retrotrapezoid nucleus neurons (underneath nVII) are positive for HA, reporter of hM4Di-HA DREADD receptor expression. The boxed area is magnified on the right. (**C-H**) Plethysmographic analysis of *Control* and *dB2-Silence* mice before (gray) and after (dark blue) CNO treatment (10 mg/kg). Breathing recordings were taken in ambient air. The precise number (n) of analyzed mice is displayed in the brackets underneath the studied ages (P7, P21 and P56). (**C**) Representative plethysmographic traces of *Control* and *dB2- Silence* mice before and after CNO. (**D**) Quantification of minute ventilation in *Control* and *dB2-Silence* mice before and after CNO. (**E, F**) Left panels, frequency distribution plots displaying tidal volumes of *Control* and *dB2-Silence* mice before and after CNO. Note that a displacement to the left indicates shallow volumes. Right panels, quantification of tidal volumes in *Control* and *dB2-Silence* mice before and after CNO. (**G, H**) Left panels, Poincaré plots illustrating breathing stability in *Control* and *dB2- Silence* mice before and after CNO. Every dot represents individual breaths, and the total number of analyzed breaths is indicated in brackets. Standard deviations (SD) 1 and 2 are displayed in Supplementary Figure 1B. Right panels, quantification of respiratory cycle lengths (T_TOT_) in control and *dB2-Silence* mice before and after CNO. Every dot in the main plots (D-H) represents the mean of individual animals. Significance was determined using one-way ANOVA followed by post hoc Tukey’s analysis.

We first verified the correct expression of the HA-tagged hM4Di receptor in a known subgroup of dB2 neurons, the retrotrapezoid nucleus, by immunostaining brainstem sections taken from *Lbx1^Cre/+^;Phox2b^FlpO/+^;RC::FPDi^+/-^* (for simplicity here called *dB2-Silence*) mice with anti-HA antibodies (Figure 1B right). Next, we compared the respiration of *dB2-Silence* mice with that of their control (*Lbx1^+/+^;Phox2b^FlpO/+^;RC::FPDi^+/-^* or *Lbx1^Cre/+^;Phox2b^+/+^;RC::FPDi^+/-^*) littermates in the absence of the synthetic DREADD agonist clozapine N-oxide (CNO) at neonatal (P7), juvenile (P21), and adult (P56) stages. No differences in minute ventilation (amount of air breathed per minute), tidal volume (amount of air taken per breath) or breathing cycle length (abbreviated here as T_Tot_), were observed between control and *dB2- Silence* mice at any of the analyzed ages (Supplementary Figure 1A). However, when the same control and *dB2-Silence* mice were treated with CNO (10 mg/kg), only the latter displayed a severe hypoventilation phenotype characterized by shallow tidal volumes that led to a diminished minute ventilation at each of the analyzed stages (Figures 1C-1F). Furthermore, the administration of CNO increased respiratory cycle lengths in *dB2-Silence* (hereafter CNO-*dB2-Silence*) neonates but had no impact on this parameter at later ages (see Figures 1G, 1H, and below). One should note that an increase in respiratory cycle length is concomitant to a reduction in breathing frequency. Interestingly, CNO-*dB2-Silence* neonates also displayed an apparent respiratory instability that was accompanied by spontaneous interruptions of breathing that ranged between 0.5 to 1.2 seconds in length, a phenotype neither observed in older CNO-*dB2-Silence* animals nor in control mice treated with CNO at any stage (Figures 1G left, 1H left, and Supplementary Figures 1B and 1C). We conclude that silencing of dB2 neuron activity has a strong effect on ventilatory control in mice and recapitulates the hypoventilation phenotype observed in CCHS patients.

### dB2 neurons are essential for the neonatal hypercarbic reflex

In humans and mice, pulmonary ventilation rapidly increases in response to elevated levels of atmospheric CO_2_ (37–39). We next evaluated whether CNO-*dB2-Silence* mice fail to respond to a hypercarbic challenge (high CO_2_ in the air; 21% O_2_, 8% CO_2_, balanced N_2_), a well-known respiratory deficit observed in CCHS patients (9, 10, 40, 41). Of note, healthy humans and mice respond to hypercarbic challenges by increasing their minute ventilation and tidal volumes, while decreasing respiratory cycle length. In the absence of CNO, both control and *dB2-Silence* mice responded similarly to the hypercarbic challenge (Supplementary Figure 2A). In contrast, CNO-*dB2-Silence* animals exhibited a severely blunted response to hypercarbia, particularly in the neonatal stage when no response to hypercarbia was observed (Figures 2A-2D). The lack of response to hypercarbia seen in CNO-*dB2-Silence* neonates was caused by a significant deficit to reduce their respiratory cycle lengths during the gas challenge (Figure 2D right). At later stages, however, CNO-*dB2-Silence* mice were partially able to reduce their respiratory cycle lengths and increase their tidal volumes during hypercarbia (Figure 2D, Supplementary Figure 2B). Thus, silencing of dB2 neuron activity precludes the onset of the hypercarbic response primarily in the neonatal period.

**Figure 2.**
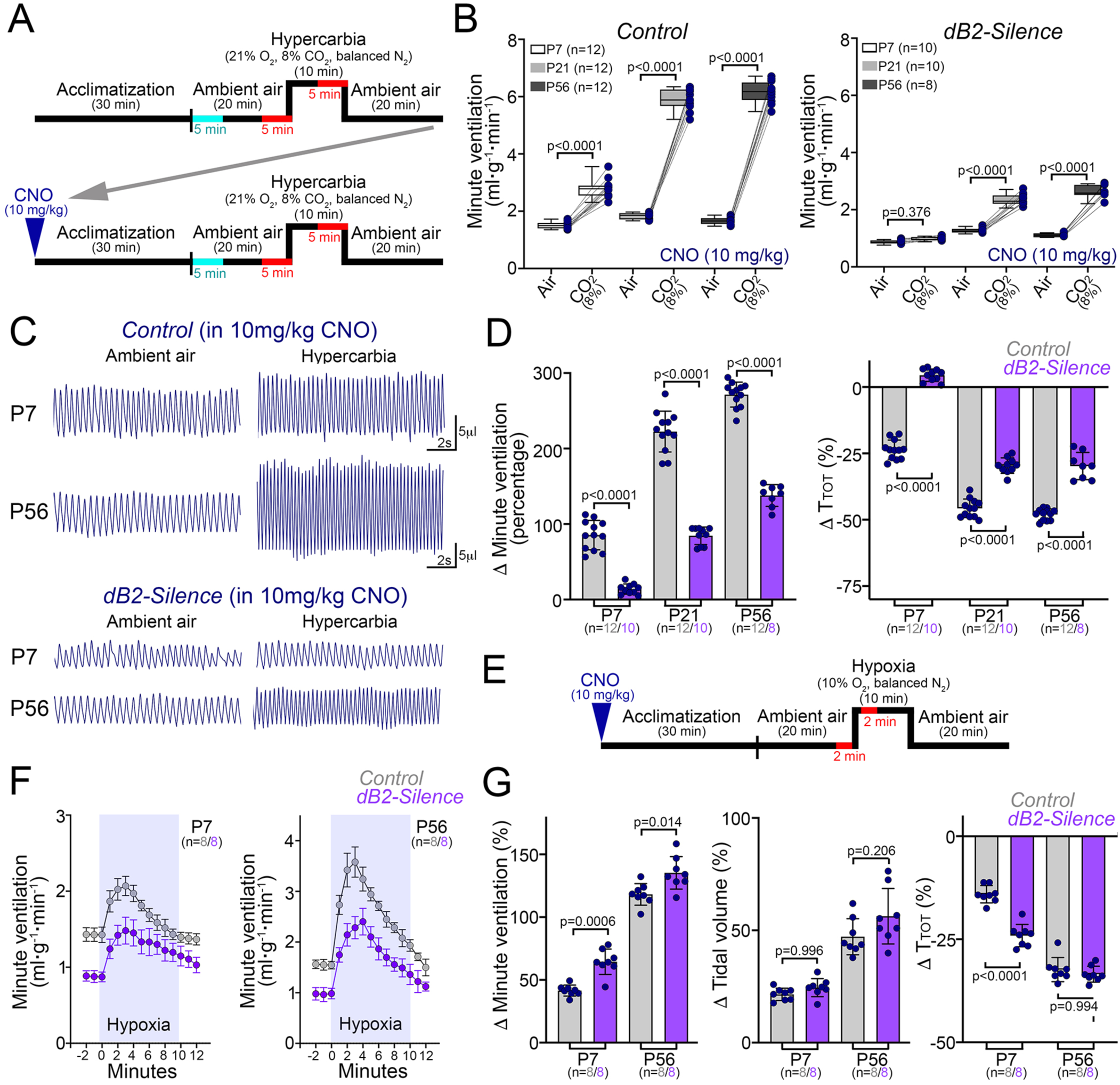
dB2 neurons are essential for neonatal hypercarbic response. (**A**) Diagram illustrating the protocol used to induce a hypercarbic response in *Control* and *dB2-Silence* mice. Respiration was analyzed in ambient air and in hypercarbia for five minutes (indicated in red). The analysis displayed in Figure 1 was obtained from the first 5 minutes (denoted in cyan) of plethysmographic recordings taken after a 30- minute acclimatization period. (**B-G**) Plethysmographic analysis of CNO-treated *Control* and *dB2-Silence* mice in ambient air (air) and in hypercarbia (8% CO_2_ in air) at the indicated ages (P7, P21 and P56). The precise number (n) of analyzed mice is displayed in brackets. Each dot in B, D and G represents the mean of individual mice. (**B**) Quantification of minute ventilation in ambient air and hypercarbia for the indicated genotypes and ages (color code: P7 white, P21 light gray, P56 dark gray). (**C**) Representative plethysmography traces of CNO-treated *Control* and *dB2-Silence* mice in ambient air and in hypercarbia. (**D**) Respiratory responses to hypercapnia expressed as percentage of change relative to the baseline (ambient air). Change (Δ) of minute ventilation (left) and respiratory cycle lengths (T_TOT_, right) for the indicated genotypes. See Supplementary Figure 2B for the quantification of tidal volumes in response to hypercarbia. (**E**) Diagram illustrating the protocol used to induce a hypoxic response in CNO-treated Control and *dB2-Silence* mice. Respiration of mice was quantified in ambient air and in hypoxia for two minutes each (denoted in red and quantified in G). (**F**) Minute ventilation values in ambient air or hypoxia (highlighted area). Each circle represents the mean ± SD over a 60-second period. (**G**) Respiratory responses to hypoxia expressed as percentage of change relative to the baseline (ambient air). Change (Δ) of minute ventilation (left), tidal volume (middle), and respiratory cycle lengths (T_TOT_, right) for the indicated genotypes. Significance was determined using one-way ANOVA followed by post hoc Tukey’s analysis.

Lastly, we assessed whether CNO-*dB2-Silence* mice were able to respond to hypoxia (low O_2_ in air; 10% O_2_, balanced N_2_; Figures 2E-2G). Despite the evident hypoventilation displayed by CNO-*dB2-Silence* mice in ambient air, no differences were observed in their ventilatory response to hypoxia when compared to CNO-treated control animals at any stage (Figures 2F, 2G). Hence, dB2 neurons are dispensable for the hypoxic reflex. Taken together, we conclude that dB2 neurons are key in the control of ventilation and specific for the regulation of the hypercarbic chemoreflex.

### Activation of dB2 neurons induces a hypercarbic-like response in neonates

To further characterize the function of dB2 neurons in respiration, we then used a second intersectional chemogenetic approach to transiently and reversibly activate these cells in neonates and adult mice. Specifically, we used the *RC::FL-hM3Dq* dual-recombinase allele that expresses an mCherry-tagged hM3Dq DREADD receptor upon the excision of two stop cassettes flanked by *FRT* and *LoxP* sites. As above, these cassettes were excised by *Lbx1^Cre^* and *Phox2b^FlpO^* (Figure 3A left). Histological analysis using *Lbx1^Cre/+^;Phox2b^FlpO/+^;RC::FL-hM3Dq^+/-^* (for simplicity *dB2-Activity*) mice confirmed the correct expression of the mCherry-tagged hM3Dq receptor in dB2 neurons, as assessed by immunoreactivity to the red fluorescent protein (to distinguish mCherry; Figure 3A right).

**Figure 3.**
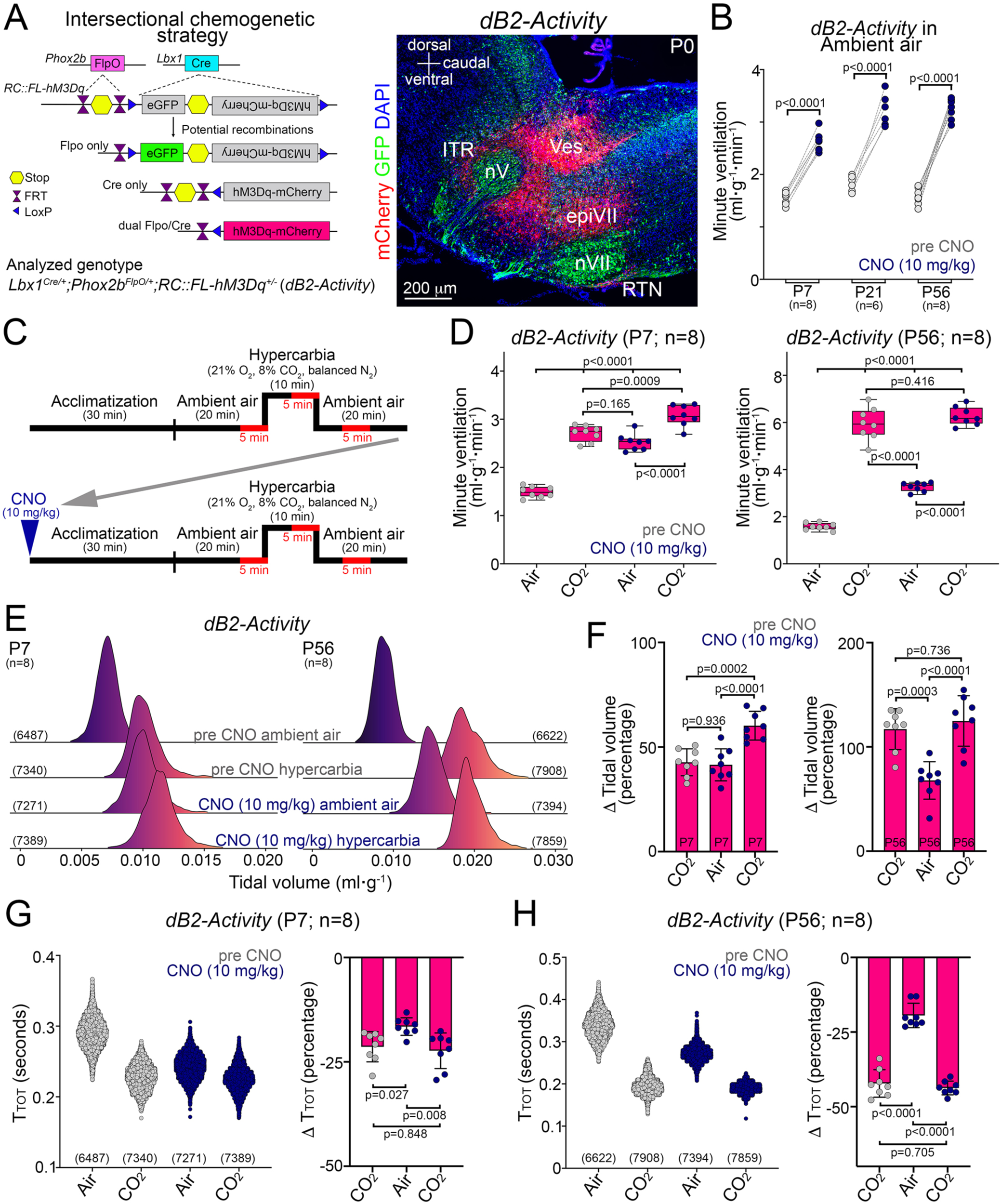
*Activation of dB2 neurons induces a full hypercarbic-like response in neonates*. (**A**) Left, intersectional genetic strategy to express the hM3Dq-mCherry DREADD receptor in dB2 neurons. The resulting *dB2-Activity* genotype is indicated. Note that FlpO-mediated recombination only allows for the expression of GFP, while the dual Cre/FlpO recombination activates hM3Dq-mCherry expression. Right, a sagittal brainstem section stained with antibodies against GFP and the red fluorescent protein (mCherry) at birth (P0). DAPI was used to counterstain. The trigeminal (nV) and facial (nVII) motor nuclei are indicated. Note that known dB2 neurons, such as intertrigeminal (ITR) and retrotrapezoid nucleus (RTN) are labeled by mCherry, reporter for hM3Dq DREADD receptor expression, but also two subgroups of newly identified dB2 neuron subgroups in this study: Ves and epiVII cells (see Figure 4 and text). (**B**) Quantification of minute ventilation in *dB2-Activity* mice before (gray) and after (dark blue) CNO treatment (10 mg/kg) at the indicated ages. The precise number (n) of analyzed mice is displayed in brackets. (**C**) Protocol used to induce a hypercarbic response in *dB2-Activity* mice at P7 and P56 (before and after CNO). (**D-H**) The respiration of *dB2-Activity* mice was analyzed in ambient air (air) and in hypercarbia (CO_2_, 8% in air) for five minutes, as indicated in red in panel C. See also Supplementary Figures 3C and 3D. (**D**) Quantification of minute ventilation of *dB2- Activity* mice in ambient air or hypercarbia (before and after CNO). (**E**) Ridge line plots illustrating the frequency distribution of tidal volumes breathed by *dB2-Activity* mice in ambient air or hypercarbia (before and after CNO). The total number of analyzed breaths is indicated in brackets. (**F**) Respiratory responses to hypercarbia expressed as percentage of change relative to the baseline (ambient air, before CNO). Change (Δ) of tidal volumes in *dB2-Activity* mice at P7 (left) and P56 (right) for the indicated conditions. (**G, H**) Left panels, scatter plots illustrating respiratory cycle length (T_TOT_) distribution in *dB2-Activity* mice at the indicated ages and conditions. The total number of analyzed breaths is indicated in brackets. Right panels, Change (Δ) of respiratory cycle lengths in response to hypercarbia in *dB2-Activity* mice at P7 and P56 for the indicated conditions. Every dot represents the mean of individual animals in D, F, G (right), H (right), or individual breaths in G (left) and H (left). Significance was determined using one-way ANOVA followed by post hoc Tukey’s analysis.

Next, we analyzed the ventilatory patterns of *dB2-Activity* mice before and after CNO administration (10 mg/kg). This showed that *dB2-Activity* mice greatly raised their ventilatory patterns when treated with CNO (CNO-*dB2-Activity*), as they increased their minute ventilation by 70%, 80%, and 110% in the neonatal, juvenile, and adult stages, respectively (Figure 3B and Supplementary Figures 3A, 3B). These augmented ventilatory patterns resulted from a pronounced increase in tidal volumes and a reduction in respiratory cycle lengths at each analyzed stage (Supplementary Figure 3B). We next compared the respiratory responses of *dB2-Activity* mice to hypercarbia before and after CNO administration at neonatal and adult stages (see experimental outline Figure 3C). In the neonatal period, untreated *dB2-Activity* mice increased their minute ventilation by 80% (relative to ambient air) and tidal volumes by 40%, while they reduced their respiratory cycle lengths by 20% when exposed to hypercarbia (Figures 3D-3H and Supplementary Figures 3C and 3D). As described above, the administration of CNO to *dB2-Activity* neonates increased their tidal volumes and reduced their respiratory cycle lengths in ambient air, changes that were virtually identical to the hypercarbic response seen in the same mice before receiving CNO (Figures 3E-3H). Thus, activation of dB2 neuron activity alone is sufficient to trigger a ventilatory response that mimics the hypercarbic reflex in neonates (Figure 3D left). Further exposure of CNO-*dB2-Activity* neonates to hypercarbia led to higher tidal volumes and minute ventilation rates than the ones observed in the same mice before CNO administration (Figures 3D-3F). In the adult period, however, CNO-*dB2- Activity* mice augmented tidal volumes and reduced respiratory cycle lengths in ambient air, but these changes represented only half the response to hypercarbia seen in the same mice before CNO administration (Figures 3E-3H). Unlike neonates, *dB2- Activity* adult mice responded to hypercarbia similarly before and after CNO (Figures 3D, Supplementary Figure 3D). We conclude that activation of dB2 neurons induces a complete hypercarbic-like response in neonates, but not in adult mice. Altogether, these data indicate that dB2 neurons are crucial for the neonatal response to hypercarbia, but only partially contribute the CO_2_ chemoreflex in the adult life.

### Spatial distribution of dB2 neurons in the brainstem

Except for the retrotrapezoid (also known as parafacial) nucleus and the intertrigeminal (or peritrigeminal) region, the location of most dB2 neurons has not been conclusively determined (12, 36, 42, 43). To define their spatial distribution, we first mapped the brainstem of mice with Lbx1 and Phox2b antibodies. Specifically, we used *Lbx1^Cre^* newborn mice carrying the reporter allele *Rosa^LSL-nGFP^* (*Lbx1^Cre/+^;Rosa^LSL-nGFP/+^* mice), a reporter that expresses a green fluorescent protein (GFP) in the nuclear membrane after Cre-mediated recombination. We used this genetic fate-mapping strategy as many Lbx1-derived neurons downregulate this factor in perinatal life (Supplementary Figure 4). This analysis uncovered five major subgroups of Lbx1+/Phox2b+ neurons that locate from cranial to caudal to: i) the intertrigeminal region, ii) the vestibular nuclei (previously defined by us as dorsal dB2 neurons; c.f 12), iii) the retrotrapezoid nucleus, iv) the dorsal part of the facial motor nucleus (here called epiVII), and v) the lateral part of the nucleus ambiguus (here called periNA) (Figure 4 and Supplementary Figure 5). We conclude that most Lbx1+/Phox2b+ (dB2) neurons locate to the rostral medulla (vestibular, retrotrapezoid and epiVII) and pons (intertrigeminal), but at least one subgroup of these cells migrates into the caudal medulla (periNA) (schematically illustrated in Figure 4D and Supplementary Figure 5E).

**Figure 4.**
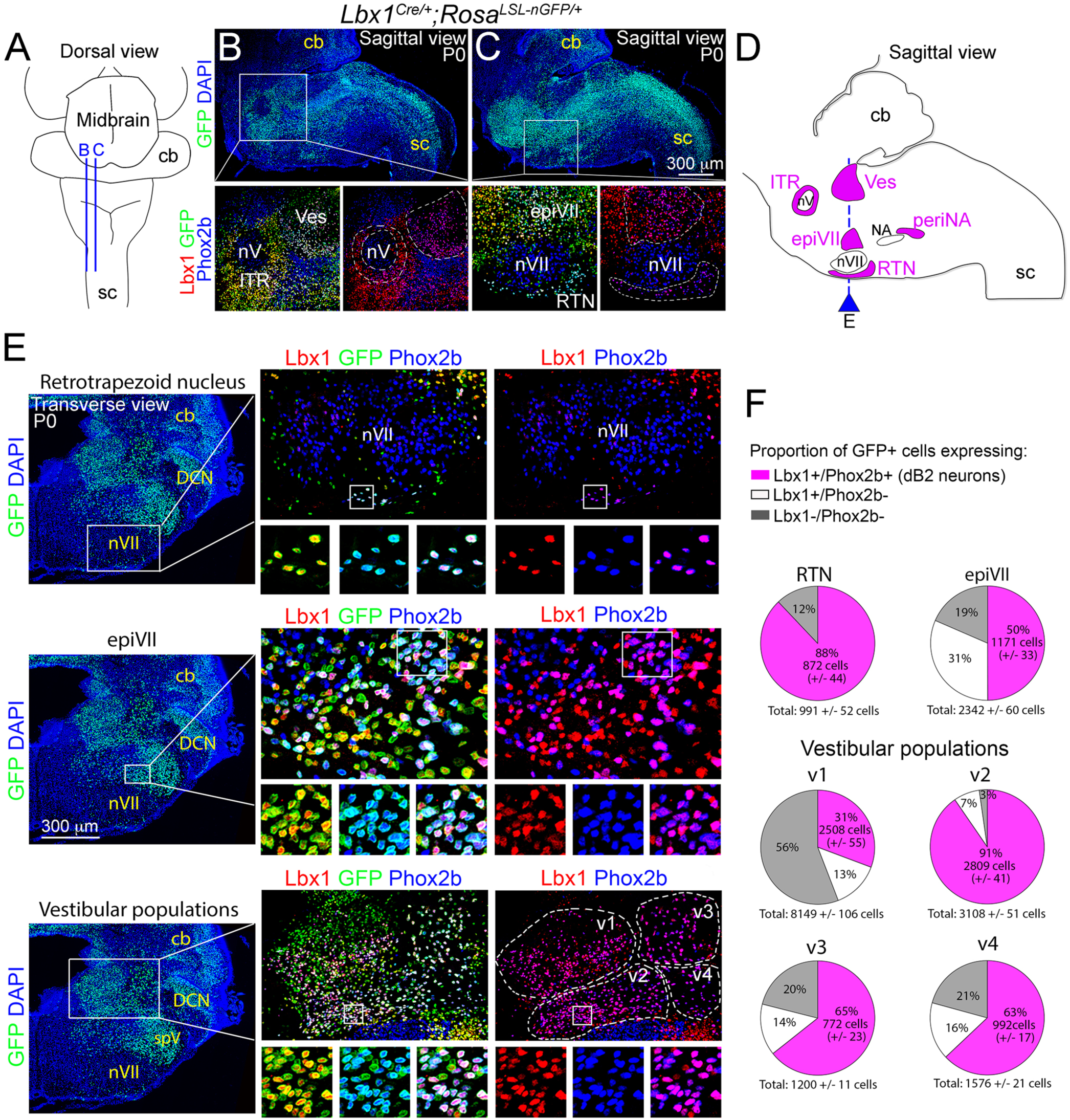
Identification of Lbx1+/Phox2b (dB2) brainstem neurons. (**A**) Schematic view of a mouse brainstem at birth (P0). Blue lines denote the sagittal planes illustrated in B and C. The spinal cord (sc) and cerebellum (cb) are displayed. (**B, C**) Sagittal sections taken from *Lbx1^cre/+^;Rosa^LSL-nGFP/+^* mice at P0. Upper panels display GFP and DAPI signals. The boxed areas are magnified at the bottom and display Lbx1, GFP and Phox2b merged signals (on the left), and Lbx1 and Phox2b only signals (on the right) for a better visualization of the Lbx1+/Phox2b+ (magenta) cells. (**D**) Schematic sagittal view of a P0 mouse brainstem illustrating the location of five Lbx1+/Phox2b+ neuron subgroups (magenta) identified here: i) intertrigeminal (ITR), vestibular (Ves), epifacial (epiVII), retrotrapezoid nucleus (RTN), and peri nucleus ambiguus (periNA) neurons. The blue line denotes the transverse plane illustrated in panel E. For analysis of ITR and periNA cells see Supplementary Figure 5. (**E**) Left, a transverse brainstem section stained with GFP and DAPI at birth. At this plane, three subgroups of Lbx1+/Phox2b+ neurons can be identified: retrotrapezoid nucleus, epiVII, and vestibular neurons. Boxed areas are magnified on the right to display Lbx1, GFP and Phox2b merged signals (middle), or Lbx1 and Phox2b only signals (right) for visualization of the Lbx1+/Phox2b+ (magenta) cells. The small boxed areas are magnified at the bottom of the main photograph. Note that four vestibular Lbx1+/Phox2b+ neuron populations can be distinguished (v1-v4), see also Supplementary Figures 5E and 6B. (**F**) Pie charts illustrating the proportion of cells with a history of Lbx1 expression (GFP+) and active expression of Lbx1 and Phox2b (dB2 neurons). The actual quantification of these cells can be found in Supplementary Figure 5D.

To reveal whether additional subgroups of dB2 neurons might exist in the brainstem, whose expression of Lbx1 and/or Phox2b becomes extinguished during their maturation, we next used a third intersectional genetic strategy to selectively mark all cells with a history of Lbx1 and Phox2b co-expression with a fluorescent reporter. To this end, we used the *RCFL-tdT* dual-recombinase allele that expresses cytoplasmatic tdTomato after the excision of two stop cassettes flanked by *FRT* and *LoxP* sites. As above, these stop cassettes were excised by *Lbx1^Cre^* and *Phox2b^FlpO^* (Figure 5A). Brains prepared from *Lbx1^Cre/+^;Phox2b^FlpO/+^;RCFL-tdT^+/-^* (for simplicity *dB2-Tomato*) newborn mice were then processed for light-sheet microscopy. 3D reconstructions of *dB2-Tomato* brains confirmed the specific location of dB2 neurons to the pons and the medulla oblongata in the five identified subgroups of Lbx1+/Phox2b+ neurons (Figures 5B, 5C, Supplementary Figure 6, and Supplementary video 1). Remarkably, this strategy also uncovered three additional subgroups of dB2 neurons in the caudal medulla who lose co-expression of Lbx1 and Phox2b by birth (Figure 5C-5E). These subgroups locate to: i) above the nucleus tractus solitarius (called here epiNTS), ii) above the nucleus ambiguus (called here epiNA), and iii) underneath the spinal trigeminal nucleus (called here infraSpV) (Figures 5D, 5E). Together, these data illustrate that although dB2 neurons retain co-expression of Lbx1 and Phox2b in the pons and rostral medulla at birth, most caudally located subgroups of dB2 neurons lose co-expression of these factors during their maturation.

**Figure 5.**
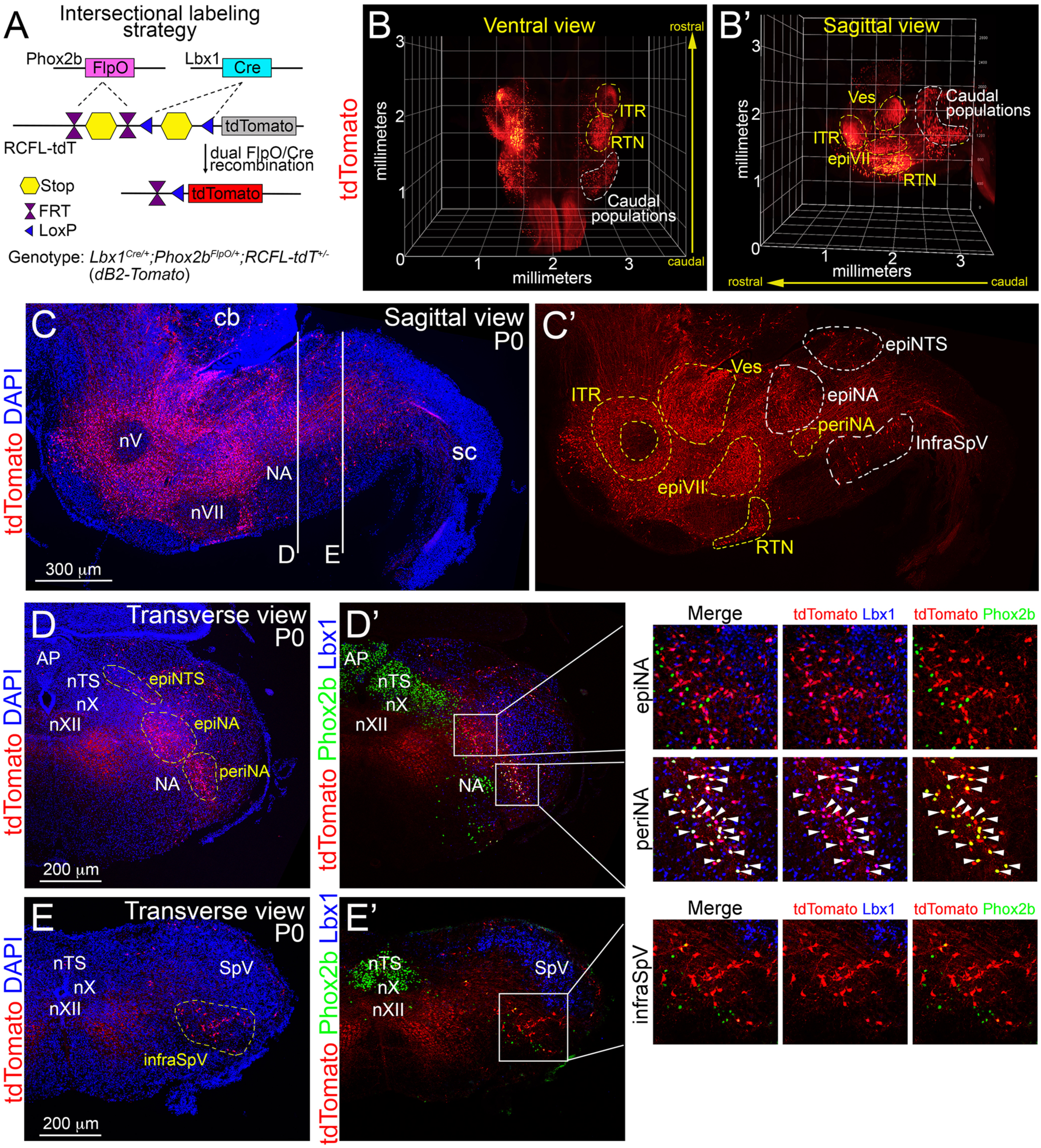
Spatial distribution of all dB2 neuron subgroups. (**A**) Intersectional genetic strategy to label with tdTomato all neurons with a history of Lbx1 and Phox2b expression, that is dB2 neurons, the resulting *dB2-Tomato* genotype is indicated. (**B, B’**) Ventral and sagittal views of 3D reconstructions created from a *dB2-Tomato* brainstem at birth (P0). tdTomato cells were distributed across the brainstem and densely packed in the intertrigeminal (ITR), vestibular (Ves), retrotrapezoid (RTN) and epifacial (epiVII) groups (yellow dashed lines). In addition, a dense group of tdTomato cells, that include peri nucleus ambiguus (periNA) cells, were seen in the caudal medulla (white dashed lines). See also Supplementary Video 1. (**C, C’**) A sagittal brainstem section stained with antibodies against the red fluorescent fluorescent protein (to detect tdTomato) and DAPI at P0. The cerebellum (cb), spinal cord (sc), trigeminal (nV), facial (nVII) and ambiguus (NA) motor nuclei are indicated for anatomical orientation. DAPI signals are removed in C’ for a better visualization of dB2 neuron subgroups. Yellow dashed lines indicate dB2 neuron subgroups that actively co-express Lbx1 and Phox2b, while white dashed lines denote dB2 neurons that lose the co-expression of these factors during their maturation. See also Supplementary Figure 6. (**D, E**) Transverse brainstem sections stained with antibodies against the red fluorescent fluorescent protein (to detect tdTomato), Lbx1 and Phox2b at P0 as indicated in C. The newly identified epiNTS, periNA, epiNA and infraSpV dB2 neurons subgroups are marked. The area postrema (AP), nucleus tractus solitarius (nTS), as well as the vagal (nX), hypoglossal (nXII) and ambiguus (NA) motor nuclei are indicated for anatomical orientation.

### dB2 neurons from rhombomeres 5 and 6 are critical for ventilation and the hypercarbic reflex

Intertrigeminal dB2 neurons originate in rhombomere 2 and are developmentally intact in homozygous *Lbx1^FS^* mice (12, 42). We therefore hypothesized that deficits in dB2 neurons generated from rhombomeres 3 to 6 might account for the respiratory phenotypes observed in CCHS patients and in homozygous *Lbx1^FS^* (ref. 12), conditional *Phox2b^Cre/+^;Lbx1^FS/lox^* (ref. 12) and chemogenetic CNO- *dB2-Silence/Activity* mice (this work). To test this, we used three distinct Cre driver mouse lines to lineage-trace the origin of distinct subgroups of dB2 neurons: *Egr2^Cre^* (expressed in rhombomeres 3 & 5), *TgHoxb1^Cre^* (expressed in rhombomere 4) and *TgHoxa3^Cre^* (expressed in rhombomeres 5 & 6) (Supplementary Figure 7). The combination of these Cre lines with the reporter *Rosa^LSL-nGFP^* line revealed the origin of most dB2 neuron subgroups that actively co-express Lbx1 and Phox2b (Supplementary Figure 8). However, caudal dB2 neurons of the periNA subgroup (and possibly other caudal dB2 populations) were not marked by any of the three Cre driver lines, suggesting either an origin of these neurons outside of rhombomeres 3 to 6 or perhaps due to the incomplete expression of Cre recombinase in rhombomere 6 by the *TgHoxa3^Cre^* mouse line (Supplementary Figure 8, and below).

We took advantage of these Cre driver lines to differentially restrict the *Lbx1^FS^* mutation to specific rhombomeres by generating *TgHoxb1^Cre/+^;Lbx1^FS/lox^* (for simplicity *r4-Lbx1^FS^*), *Egr2^Cre/+^;Lbx1^FS/lox^* (*r3&5-Lbx1^FS^*), and *TgHoxa3^Cre/+^;Lbx1^FS/lox^* (*r5&6- Lbx1^FS^*) mice. These conditional animals were then analyzed by head-out plethysmography immediately at birth. As compared to littermate controls, *r4-Lbx1^FS^* pups displayed no significant differences in their minute ventilation, tidal volumes nor in their respiratory cycle lengths while breathing ambient air (Figures 6A-6D). In contrast, *r3&5-Lbx1^FS^* newborns showed a mild hypoventilation phenotype characterized by longer respiratory cycle lengths but no significant change in their tidal volumes (Figures 6A-6D). Strikingly, *r5&6-Lbx1^FS^* newborns displayed a severe hypoventilation phenotype that resulted from shallow tidal volumes, aberrantly long respiratory cycle lengths, and frequent apneic events that lasted from 3 to 30 seconds (Figures 6A-6D and Supplementary Figure 9A). Notably, the respiratory phenotype observed in *r5&6-Lbx1^FS^* newborns closely resembled to that reported for homozygous *Lbx1^FS^* and conditional *Phox2b^Cre/+^;Lbx1^FS/lox^* animals (12).

**Figure 6.**
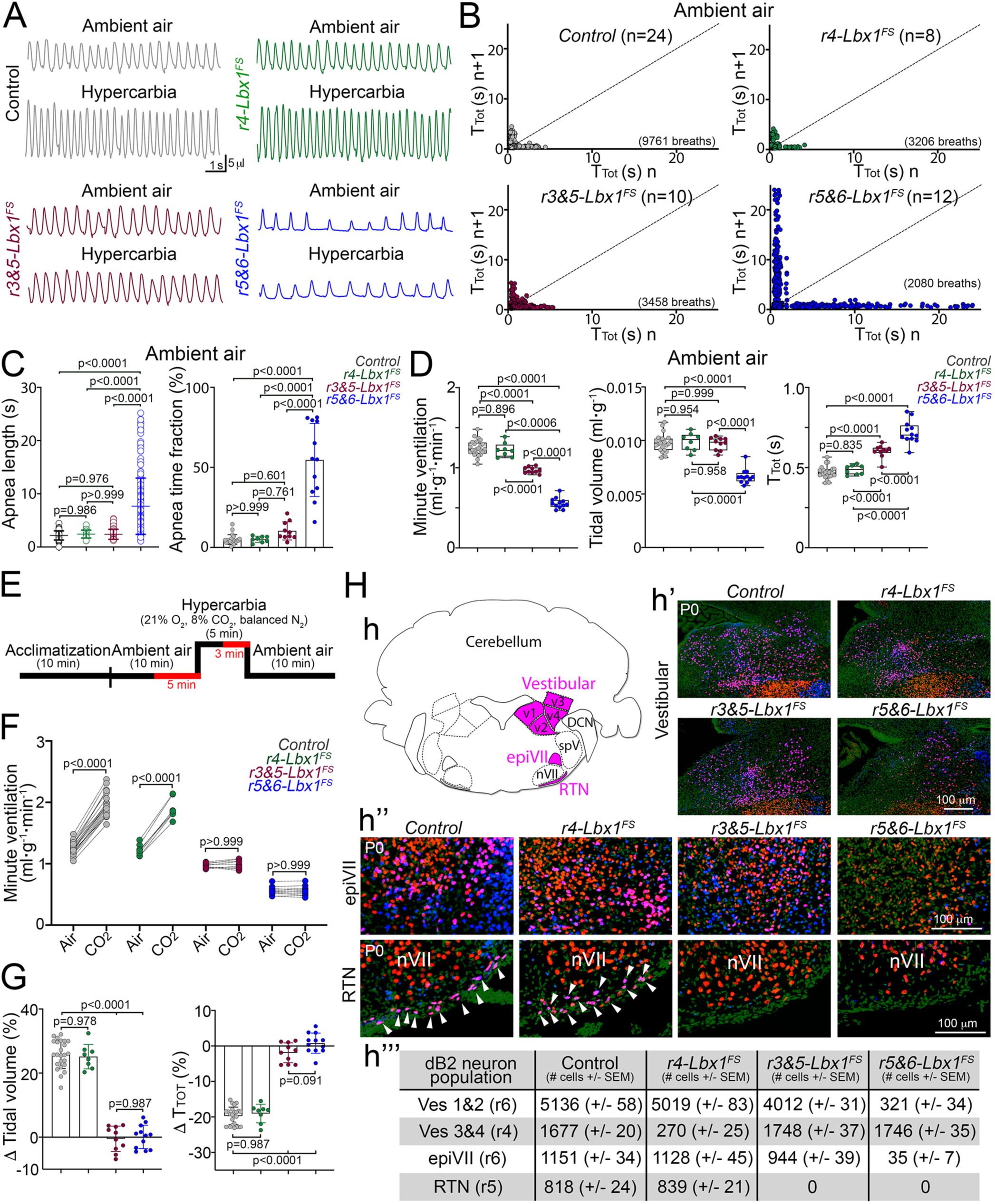
dB2 neurons originated from rhombomeres 5 and 6 are essential for ventilatory control and the hypercarbic reflex at birth. Plethysmographic and anatomical analyses of *Control*, *(Tg)Hoxb1^Cre/+^;Lbx1^FS/lox^* (*r4-Lbx1^FS^*), *Egr2^cre/+^;Lbx1^FS/lox^* (*r3&5-Lbx1^FS^*), and *(Tg)Hoxa3^cre/+^;Lbx1^FS/lox^* (*r5&6-Lbx1^FS^*) newborn (P0) mice. (**A**) Representative plethysmography recordings of *Control*, *r4-Lbx1^FS^*, *r3&5-Lbx1^FS^*, and *r5&6-Lbx1^FS^* newborns in ambient air and hypercarbia. (**B**) Poincaré plots of breathing stability of *Control*, *r4-Lbx1^FS^*, *r3&5-Lbx1^FS^*, and *r5&6-Lbx1^FS^* newborns while breathing ambient air. Every dot represents individual breaths. Quantification of SD1 and SD2 can be found in Supplementary Figure 9A. The number of analyzed breaths and (n) mice are indicated in brackets. (**C**) Quantification of apnea lengths (left) and the fraction of time in apnea (right) of *Control*, *r4-Lbx1^FS^*, *r3&5- Lbx1^FS^*, and *r5&6-Lbx1^FS^* newborns while breathing ambient air. Each circle represents individual apneas (left), while each dot the mean of individual mice (right). (**D**) Quantification of minute ventilation, tidal volume, and respiratory cycle lengths (T_TOT_) in *Control*, *r4-Lbx1^FS^*, *r3&5-Lbx1^FS^*, and *r5&6-Lbx1^FS^* newborns while breathing ambient air. (**E**) Diagram illustrating the protocol used to induce a hypercarbic response in newborn mice. The respiration of mice was analyzed in ambient air (5 minutes) and in hypercarbia (3 minutes) as indicated (in red). (**F**) Quantification of minute ventilation of *Control*, *r4-Lbx1^FS^*, *r3&5-Lbx1^FS^*, and *r5&6-Lbx1^FS^* newborns while breathing ambient air or hypercarbia. (**G**) Respiratory responses to hypercarbia expressed as percentage of change relative to the baseline (ambient air). Change (Δ) of tidal volume (left) and respiratory cycle lengths (right) displayed by the indicated genotypes. (**H**) Histological characterization of *Control*, *r4-Lbx1^FS^*, *r3&5-Lbx1^FS^*, and *r5&6-Lbx1^FS^* newborns. (**h**) Schema illustrating the location of vestibular (v1-v4), epifacial (epiVII) and retrotrapezoid nucleus (RTN) dB2 neurons. (**h’-h’’**) Histological analyses of the indicated genotypes using antibodies against Lbx1 and Phox2b. DAPI (in green) was used to counterstain. Co-expression of Lbx1 and Phox2b results in a magenta coloration. See also Supplementary Figure 10A. (**h’’’**) Quantification summary of the indicated dB2 neuron subgroups and genotypes. See Supplementary Figures 10B and 10C for the actual quantification of these dB2 neuron subgroups. Each dot represents the mean of individual mice in D, F and G. Significance was determined using one-way ANOVA followed by post hoc Tukey’s analysis.

Next, we evaluated the response of *r4-Lbx1^FS^*, *r3&5-Lbx1^FS^* and *r5&6-Lbx1^FS^* newborns to hypercarbia. This demonstrated that *r4-Lbx1^FS^* pups efficiently respond to the chemochallenge. In contrast, neither *r3&5-Lbx1^FS^* nor *r5&6-Lbx1^FS^* newborns showed a response to the gas exposure (Figures 6E-6G). Lastly, we used immunofluorescence to examine the brainstems of *r4-Lbx1^FS^*, *r3&5-Lbx1^FS^* and *r5&6- Lbx1^FS^* newborns to determine any potential changes in the dB2 neuron subgroups described in Figure 4. This illustrated the unique loss of lateral vestibular (v3 and v4 subgroups) dB2 neurons in *r4-Lbx1^FS^* pups; the single loss of retrotrapezoid nucleus neurons in *r3&5-Lbx1^FS^* newborns; and the loss of medial vestibular (v1 and v2 subgroups) dB2 neurons, epiVII cells and retrotrapezoid nucleus neurons in *r5&6- Lbx1^FS^* animals (Figure 6H, quantified in Supplementary Figures 10A-10D). One should note that the periNA population was unchanged in each of the three analyzed genotypes, suggesting that caudal dB2 neuron subgroups might be unaffected in *r4- Lbx1^FS^*, *r3&5-Lbx1^FS^* and *r5&6-Lbx1^FS^* animals (Supplementary Figure 10E, and below). Taking together these data, we conclude that: i) lateral vestibular dB2 neurons are dispensable for breathing, ii) loss of retrotrapezoid nucleus neurons in *r3&5-Lbx1^FS^* and in *r5&6-Lbx1^FS^* newborns correlates with prolonged respiratory cycle lengths and the inability to respond to hypercarbia, and iii) the loss of medial vestibular dB2 neurons and epiVII cells in *r5&6-Lbx1^FS^* newborns correlates with shallow tidal volumes and the appearance of apneic behavior.

Unlike homozygous *Lbx1^FS^* and conditional *Phox2b^Cre/+^;Lbx1^FS/lox^* animals (12), all conditional *r4-Lbx1^FS^*, *r3&5-Lbx1^FS^* and *r5&6-Lbx1^FS^* pups survive the perinatal period. This allowed us for the first time to analyze their breathing behavior at later postnatal stages: P7, P21 and P56. As in the newborn period, *r4-Lbx1^FS^* mice did not display any obvious impairment either in ventilation or in response to hypercarbia at any of the analyzed postnatal stages (Figure 7A and Supplementary Figures 11A, 11B). This confirmed that dB2 neuron subgroups derived from the rhombomere 4 are dispensable for breathing. Plethysmographic recordings of *r3&5-Lbx1^FS^* animals still showed a mild hypoventilation at P7, but no ventilatory deficit at more mature (P21 or P56) stages while breathing ambient air (Figure 7A and Supplementary Figure 11A). In contrast, *r5&6-Lbx1^FS^* mice significantly hypoventilate throughout postnatal life in ambient air (Figure 7A and Supplementary Figure 11A). This deficit was mainly due to shallow tidal volumes produced by *r5&6-Lbx1^FS^* animals (Supplementary Figure 11A). Interestingly, respiratory cycle lengths were only partially increased in both *r3&5- Lbx1^FS^* and *r5&6-Lbx1^FS^* neonates but did not differ from control animals at more mature stages (Supplementary Figure 11A). Furthermore, the incidence of apneic events in *r5&6-Lbx1^FS^* newborn mice did not significantly differ to control animals (or to *r4-Lbx1^FS^* or *r3&5-Lbx1^FS^* mice) after the neonatal period (Supplementary Figure 9B). Together, these data show that elimination of medial vestibular dB2 neurons and epiVII cells (in *r5&6-Lbx1^FS^* mice) results in shallow tidal volumes throughout postnatal life, while the absence of retrotrapezoid nucleus neurons (in *r3&5-Lbx1^FS^* and *r5&6- Lbx1^FS^* mice) affects respiratory cycle lengths specifically in neonates.

**Figure 7.**
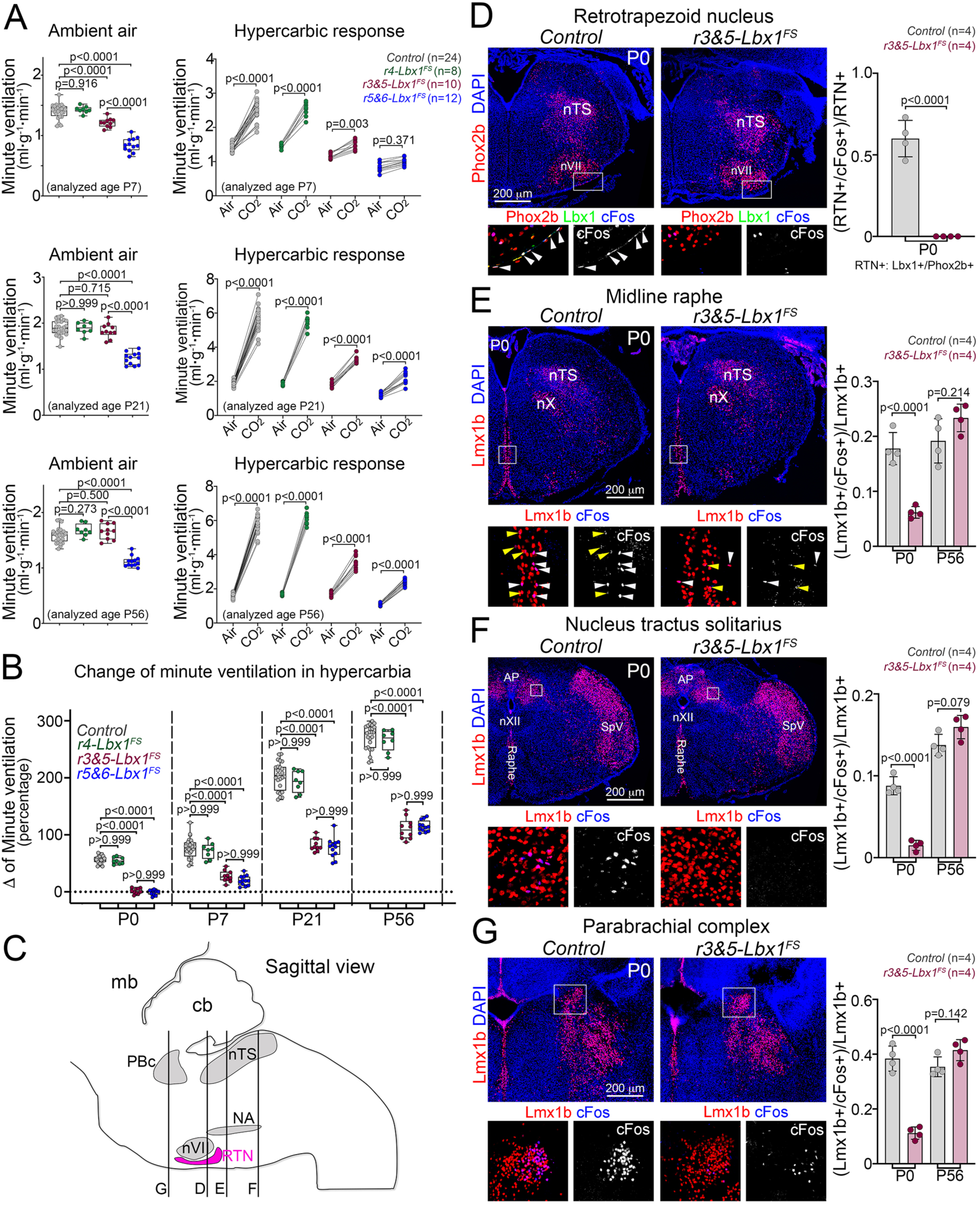
Partial recovery of the hypercarbic reflex in mature *r3&5-Lbx1^FS^* and *r5&6-Lbx1^FS^* mice. (**A**) Quantification of minute ventilation displayed by *Control*, *r4- Lbx1^FS^*, *r3&5-Lbx1^FS^*, and *r5&6-Lbx1^FS^* mice at the indicated ages while breathing ambient air (left panels) or their response to hypercarbia (right panels). The (n) number of analyzed mice is displayed in brackets. See also Supplementary Figures 11A. (**B**) Respiratory response to hypercarbia expressed as percentage of change relative to the baseline (ambient air). Change (Δ) of minute ventilation display by the indicated genotypes at different postnatal ages: P0, P7, P21 and P56. See also Supplementary Figure 11B. (**C**) Schematic sagittal view of a mouse brainstem displaying the location of the transverse planes shown in panels D-G. (**D-G**) Left panels, histological detection of cFos+ cells after 1 hour exposure to hypercarbia in *Control* and *r3&5-Lbx1^FS^* mice at birth (P0). The nucleus tractus solitarius (nTS), area postrema (AP), as well as the facial (nVII), vagal (nX), and hypoglossal (nXII) motor nuclei are indicated for anatomical orientation. Retrotrapezoid nucleus neurons were detected with Lbx1 and Phox2b antibodies (D). Raphe (E), nucleus tractus solitarius (F), and parabrachial complex (G), neurons were detected with Lmx1b antibodies. Boxed areas in the main photographs are magnified at the bottom. Magnifications display merged or cFos only signals. White and yellow arrowheads in F denote neurons with strong and weak cFos immunoreactivity, respectively. See Supplementary Figure 11C for the histological analysis of *r3&5-Lbx1^FS^* at P56. Right panels, quantification of retrotrapezoid nucleus, nucleus tractus solitarius, middle raphe, and parabrachial complex neurons co-expressing cFos at P0 and P56. Each dot represents the mean of individual mice. Significance was determined using one-way ANOVA followed by post hoc Tukey’s analysis.

Next, we analyzed the ventilatory responses of *r3&5-Lbx1^FS^* and *r5&6-Lbx1^FS^* animals to hypercarbia at P7, P21 and P56. In contrast to the lack of response to the gas challenge seen in both genotypes at birth, we detected a progressive increase in their ventilatory responses to the chemochallenge from the neonatal stage (P7) to the adulthood (P56) (Figure 7B). One should note, however, that the maximal response to hypercarbia seen in adult *r3&5-Lbx1^FS^* and *r5&6-Lbx1^FS^* animals represented only 1/3 of that observed in control mice of the same age (Figures 7A, 7B and Supplementary Figure 11B). Thus, retrotrapezoid nucleus neurons are crucial for the hypercarbic response during the neonatal life, but the response to hypercarbia appears to result from the cooperation of retrotrapezoid nucleus neurons with additional chemoreceptor neurons as the animal matures.

To understand how the loss of retrotrapezoid nucleus neurons affects the central circuit mediating the hypercarbic reflex, we stained brainstem sections taken from newborn and adult *r3&5-Lbx1^FS^* mice, after an hour-long exposure to hypercarbia, with antibodies against cFos to label neurons activated by the chemochallenge (Figures 7C-7G). We analyzed *r3&5-Lbx1^FS^* animals for this experiment as they specifically and uniquely lack retrotrapezoid nucleus neurons. As expected, cFos+ cells were undetected in the ‘retrotrapezoid nucleus’ of *r3&5-Lbx1^FS^* mice (Figure 7D). Surprisingly, we hardly detected cFos+ cells in other regions known to participate in the hypercarbic reflex, such as the midline medullary raphe (that is, raphe obscurus, magnus and pallidus), the caudal part of the nucleus tractus solitarius or the parabrachial complex in *r3&5-Lbx1^FS^* newborns (Figure 7E-7G). In contrast, no difference in the number of cFos+ cells was observed in the midline raphe, nucleus tractus solitarius and parabrachial complex in mature *r3&5-Lbx1^FS^* animals when compared to controls of the same age (Figure 7E-7G and Supplementary Figure 11C). We conclude that retrotrapezoid nucleus neurons are a prerequisite for the activation of the central circuit mediating hypercarbia at birth, but most elements of this circuit are activated independently of the retrotrapezoid nucleus in the adult life.

### Loss of caudal dB2 neurons leads to perinatal lethality

Despite the close phenotypic resemblance of *r5&6-Lbx1^FS^* newborns with homozygous *Lbx1^FS^* and conditional *Phox2b^Cre/+^;Lbx1^FS/lox^* pups in terms of hypoventilation patterns, shallow tidal volumes and frequent apneic behavior, no *r5&6-Lbx1^FS^* animal died at birth. We then asked whether periNA cells, as well as the other previously unidentified caudal dB2 subgroups (epiNA, epiNTS and infraSpV), were affected in the *Lbx1^FS^* background. To this end, we extended the use of our intersectional *dB2-Tomato* line to bear the *Lbx1^FS^* mutation and determined changes in these caudal populations by immunofluorescence. This showed that all dB2 caudal subgroups were mis-specified and underwent marked fate-shifts in *Lbx1^Cre/FS^;Phox2b^Flpo/+^;RCFL-tdT^+/-^* (for simplicity *dB2-Tomato-Lbx1^FS^*) mice (Supplementary Figure 12). For instance, cells of the epiNTS subgroup were converted into Phox2b+ neurons of the nucleus tractus solitarius, whereas the periNA, epiNA, and infraSpV subgroups appeared to be mislocated to the spinal somatosensory trigeminal nucleus (Supplementary Figure 12). We conclude that loss of caudal dB2 neurons in homozygous *Lbx1^FS^* newborns might cause the lethality of the genotype.

We next compared the recombination pattern of the *TgHoxa3^Cre^* line with a second Cre driver line that has also been reported to recombine rhombomeres 5 and 6 during early development, *MafB^Cre^* (44). Specifically, we compared the recombination patterns of these two Cre lines at E11.5, the developmental timepoint when dB2 neurons are specified (12, 35). To accurately map the borders of these rhombomeres, we also incorporated the *Egr2^Cre/+^* driver line into our analysis. Cre recombination was visualized with the *Rosa^LSL-nGFP^* reporter. This revealed that *MafB^Cre^,* but not *TgHoxa3^Cre^*, recombines the full extension of rhombomere 6 (Supplementary Figure 13). Next, we used the *MafB^Cre^* driver line to restrict the *Lbx1^FS^* mutation to rhombomeres 5 and 6 and generated *MafB^Cre/+;^Lbx1^FS/lox^* (for simplicity *MafB-Lbx1^FS^*) animals. Remarkably, 4/12 *MafB-Lbx1^FS^* newborns immediately died at birth without any apparent breathing behavior. The remaining (8/12) rarely displayed a continuous breathing pattern, but instead exhibited notable gasping behavior with prolonged apneic times and survived for a maximum of 2 hours after delivery (Figures 8A-8C and Supplementary Video 2). Furthermore, *MafB-Lbx1^FS^* newborns did not display a hypercarbic response (Supplementary Figures 14A, 14B). Histological examination of *MafB-Lbx1^FS^* pups using Lbx1 and Phox2b antibodies revealed the absence of a recognizable periNA group, in addition to the loss of medial vestibular dB2 neurons, epiVII cells and retrotrapezoid nucleus neurons (Figure 8D and Supplementary Figures 14C, 14D). We conclude that dB2 neurons originating from rhombomere 6 are required for ventilatory control, correct tidal volumes and neonatal survival (Summarized in Figures 8E-8G).

**Figure 8.**
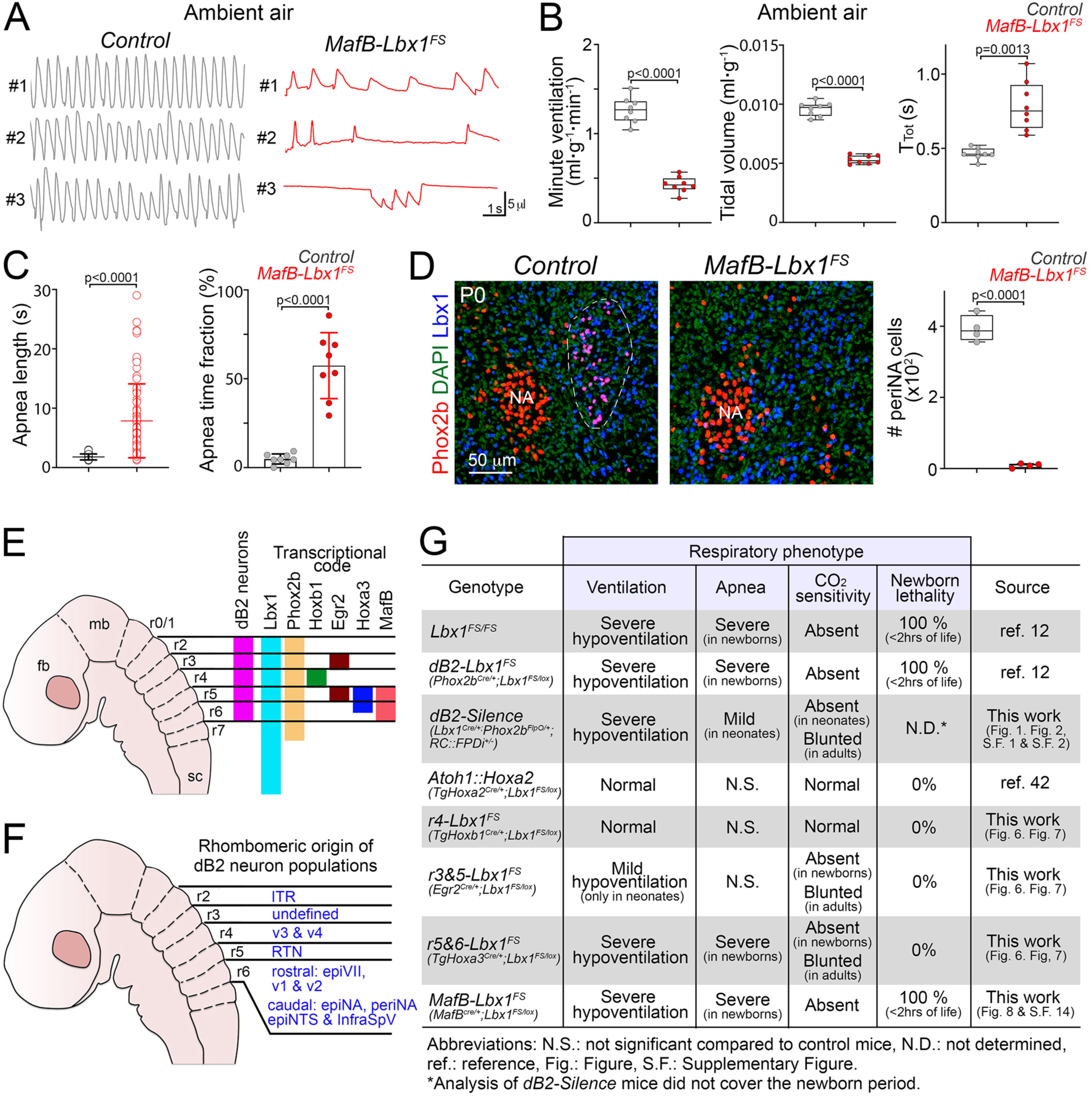
Loss of caudal dB2 neurons causes respiratory arrest at birth. (**A**) Representative plethysmography traces of three distinct *Control* and three different *MafB^cre/+^;Lbx1^FS/lox^ (MafB-Lbx1^FS^*) newborn mice while breathing ambient air. (**B**) Quantification of minute ventilation, tidal volume, and respiratory cycle lengths (T_TOT_) in *Control* and *MafB-Lbx1^FS^* newborns while breathing ambient air. Each dot represents the mean of individual mice. (**C**) Quantification of apnea lengths (left) and the fraction of time in apnea (right) of *Control* and *MafB-Lbx1^FS^* newborn mice while breathing ambient air. Each circle represents individual apneas (left), while each dot the mean of individual mice (right). (**D**) Histological analysis and quantification of peri nucleus ambiguus (periNA) neurons in *Control* and *MafB-Lbx1^FS^* newborns at birth (P0). The transverse brainstem sections were stained with Lbx1 and Phox2b antibodies. DAPI (in green) was used to counterstain. See Supplementary Figure 14 for the analysis of other dB2 neuron subgroups in *MafB-Lbx1^FS^* mutants. (**E**) Schema displaying the rhombomeric (r) segmentation of the brainstem, the rhombomeric origin of dB2 neurons, and expression patterns of the genes analyzed in this study. The forebrain (fb), midbrain (mb) and spinal cord (sc) are indicated for anatomical orientation. (**F**) Schema summarizing the rhombomeric origin of the identified dB2 neuron subgroups in this study. (**G**) Summary of the main findings of this study. The phenotypes of the previously reported homozygous *Lbx1^FS^* (ref. 12), conditional *Phox2b^Cre/+^;Lbx1^FS/lox^* (ref. 12) and *Atoh1::Hoxa2* (*TgHoxa2^Cre/+^;Atho1^LacZ/lox^*, ref. 42) are also displayed for comparison. Two-tailed t-tests was performed to determine statistical significance in B-D.

## Discussion

In this study, we investigated the function of medullary dB2 neurons in ventilatory control and their relevance to CCHS. Our findings demonstrate that these neurons are critical for respiratory tidal volumes, as well as key for neonatal respiratory stability, neonatal respiratory frequency patterns and the hypercarbic reflex. Our intersectional studies and conditional mutagenesis identified a single subgroup of dB2 neurons (generated from rhombomere 5), as an obligatory element for the central circuit regulating the neonatal hypercarbic reflex and neonatal respiratory frequencies. These experiments also uncovered novel components of the neural circuit serving in ventilatory control (generated from rhombomere 6), whose nullification results in various manifestations of the CCHS phenotype: i) hypoventilation due to shallow tidal volumes, ii) respiratory instability, and iii) respiratory arrest.

In previous work, we identified a recessive mutation in *LBX1* (*LBX1^FS^*) to be causative of severe CCHS in children (12). Mice bearing an analogous mutation die immediately at birth from an apparent failure to breathe and present with a unique deficit in the development of two dB2 neuron subgroups, the retrotrapezoid nucleus and a group of dorsally located cells that we now define as vestibular dB2 (v1-v4) neurons (ref. 12 and this work). However, the sudden and premature death of homozygous *Lbx1^FS^* newborn mice hampered the physiological association of dB2 neurons with the breathing phenotype. In this study, we now produced new murine models for the specific study of dB2 neurons and report that their selective invalidation results in severe hypoventilation, diminished tidal volumes, slow neonatal respiratory frequencies, and an anomalous hypercapnic reflex, which collectively recapitulate CCHS in mice. Our findings are of both clinical and biological relevance. From a clinical perspective, this work establishes dB2 neuron dysfunction to be causative of CCHS and provides the missing link between this disorder and the known genetic perturbations (*PHOX2B* and *LBX1*) that affect CCHS patients. Furthermore, our new murine models also offer the opportunity for the development of future therapeutic strategies for the management of CCHS, without the confounding factor of autonomous nervous system dysregulation seen in other models, such as mice carrying *PHOX2B* mutations. From the biological standpoint, our work uncovers novel neuronal components of the respiratory circuit regulating homeostatic breathing.

The shallow respiratory tidal volumes seen in CNO-*dB2-Silence* mice, while breathing ambient air, are attributable to dB2 neurons located either in the medial vestibular nuclei (v1 and v2 subgroups) and/or above the facial motor nucleus (epiVII subgroup). Our systematic lineage-tracing studies show that these neurons emerge from rhombomere 6 and their absence in *r5&6-Lbx1FS* mice causes comparable deficits in tidal volumes to the ones observed in CNO-*dB2-Silence* animals and in CCHS. Early studies showed that electrical or chemical stimulation to activate or inhibit medial vestibular nuclei, or their afferents (cerebellar fastigial cells), can induce respiratory responses that augment or diminish ventilatory patterns via prominent effects on tidal volumes, and that these changes are lost or attenuated by the bilateral destruction of the medial vestibular nuclei (45–49). Therefore, it is tempting to speculate that the diminished tidal volumes observed in CNO-*dB2-Silence* and *r5&6- Lbx1^FS^* mice might be a direct result of the inactivation and absence of medial vestibular dB2 neurons, respectively. Nonetheless, further work is necessary to exclude the newly identified epiVII group from tidal volume control.

Silencing of dB2 neuron activity abrogates the hypercarbic chemoreflex in neonates and severely blunts it in juvenile and adult mice. The dB2 neuron subgroup responsible for this is the retrotrapezoid nucleus. Indeed, *r3&5-Lbx1^FS^* and *r5&6- Lbx1^FS^* newborns, which lack retrotrapezoid nucleus neurons, are fully insensitive to hypercarbia at birth. The retrotrapezoid nucleus has long been considered as a major node for sensing changes of pCO_2_ within the central nervous system (50, 51). Here, we show that the lack of these neurons precludes the activation of other neurons associated with the hypercarbic response, such as the midline raphe (raphe obscurus, magnus and pallidus), the nucleus tractus solitarius, and the parabrachial complex, at birth. Given that retrotrapezoid nucleus neurons form reciprocal connections with these brainstem centers (38, 52), our data indicate that retrotrapezoid nucleus neurons are not only a central node, but are obligatory for the efficient activation of the respiratory circuit driving the response to hypercarbia in neonates. Although severely blunted, a significant hypercarbic response was detected in juvenile and adult *r3&5- Lbx1^FS^* and *r5&6-Lbx1^FS^* animals. In keeping with this, we observed activation of midline raphe, nucleus tractus solitarius and parabrachial cells in mature *r3&5-Lbx1^FS^* animals following a hypercarbic exposure. These data imply that raphe, nucleus tractus solitarius and parabrachial cells can be activated independently of the retrotrapezoid nucleus to modulate part of the hypercarbic reflex in mature life.

We show here that the loss of retrotrapezoid nucleus neurons, in *r3&5-Lbx1^FS^* and *r5&6-Lbx1^FS^* mice, resulted in increased respiratory cycle lengths only in the neonatal period. The same increase in respiratory cycle lengths can be observed in CNO-*dB2-Silence* neonates, but not in more mature animals. Although with different degrees of technical specificity, the optogenetic activation of retrotrapezoid nucleus neurons, via viral transductions, can increase baseline respiratory frequencies and tidal volumes in adult mice (53, 54). However, we show here that neither juvenile nor adult *r3&5-Lbx1^FS^* mice, whose only recognizable deficit is the absence of retrotrapezoid nucleus neurons, display any significant differences in baseline tidal volumes nor in respiratory cycle lengths when compared to control animals. This reveals that retrotrapezoid nucleus neurons have no function in baseline respiration after the neonatal period. Even though not discussed in detail, a similar result was obtained by Ramanantsoa et al., (2011) using a mouse model that lacks a substantial number of retrotrapezoid nucleus neurons (28). Nonetheless, the activation of retrotrapezoid nucleus neurons in our CNO-*dB2-Activity* mice led to augmented baseline tidal volumes and respiratory frequencies from neonates to adults, but these effects most likely represent the induction of a hypercarbic-like response. In support of this, *dB2-Activity* neonates display identical changes in minute ventilation, tidal volumes, and respiratory cycle lengths after CNO, to the ones observed in the same mice before CNO in hypercarbia. In the adult period, CNO-*dB2-Activity* mice also augmented tidal volumes and reduced respiratory cycle lengths in ambient air, but these changes were only half the response to hypercarbia seen in the same mice before CNO administration. Our data thus reveal that retrotrapezoid neurons are essential for the hypercarbic reflex and the control of respiratory frequencies in neonatal life, but only contribute to hypercarbia in adults.

The severe apneic phenotype previously seen in homozygous *Lbx1^FS^* pups (12), and now here in *r5&6-Lbx1^FS^* and *MafB-Lbx1^FS^* newborns is not attributable to the lack of retrotrapezoid nucleus neurons, as no obvious change in the incidence of apneas was seen in *r3&5-Lbx1^FS^* newborns when compared to control littermates. On the contrary, this apneic phenotype correlated with the absence of dB2 neurons in either the medial vestibular nuclei or the epiVII region seen in *r5&6-Lbx1^FS^* and *MafB- Lbx1^FS^* newborns. Since no significant apneic behavior was reported after inhibition or destruction of the medial vestibular nuclei (46, 49), we hypothesize that the newly identified epiVII subgroup might have an anti-apneic function in neonates. A word of caution here is that previous work that inhibited or destroyed the medial vestibular nuclei was conducted in adult animals, not in neonates, and we did not detect any significant apneic behavior in *r5&6-Lbx1^FS^* nor in CNO-treated *dB2-Silence* adult animals.

The sudden respiratory arrest seen in homozygous *Lbx1^FS^* pups (12) and reproduced here in *MafB-Lbx1^FS^* newborns seems to be attributable to the compound loss of caudally (rhombomere 6) generated dB2 neuron groups. These include the periNA, epiNA, epiNTS, and infraSpV, in addition to medial vestibular dB2 (v1 and v2) subgroups and epiVII cells. We show that these neurons undergo marked fate- changes during their development that prevent their correct specification. Future work is necessary to assess the potential function of these previously unknown populations in homeostatic respiration, as well as to define whether these cells individually or synergistically support neonatal survival.

In summary, our work identifies specific subgroups of medullary dB2 neurons that coalesce into novel brainstem centers regulating breathing homeostasis. These neurons originate from rhombomeres 5 and 6 and directly participate in ventilatory control, respiratory stability and the hypercarbic reflex. We also show that other dB2 neurons (from rhombomeres 3 and 4) are not necessary for respiration. Although intertrigeminal dB2 neurons, a rhombomere 2 derivative, are developmentally unaffected in homozygous *Lbx1^FS^* mice and not targeted in *r5&6-Lbx1^FS^* nor in *MafB- Lbx1^FS^* newborns (this work and ref. 12), available evidence indicates that these cells might also play a role in homeostatic respiration and partially on the accuracy of tidal volumes (42). Thus, multiple dB2 neuron subgroups are hard-wired to the central respiratory circuit. Collectively, our studies establish dB2 neuron dysfunction to be causative of hypoventilation and other respiratory phenotypes characteristically seen in CCHS.

### Animals

All animal experimental procedures were approved by the Landesamt für Gesundheit und Soziales (G-0002/21) and the Charité Universitätsmedizin Berlin (TCH-0008/21). Mouse lines used in this study: *Egr2^Cre^* (55), *Lbx1^Cre^* (56), *Lbx1^lox^* (12), *Lbx1^FS^* (12), *MafB^Cre^* (57), *Phox2b^FlpO^* (58), *TgHoxb1^Cre^* (59, 60), *TgHoxa3^Cre^* (61), *RC::FPDi* (62), *RC::FL-hM3Dq*, (63), *Rosa^LSL-nGFP^* (64) and *RCFL-tdT^+/-^* (65). All mouse lines were maintained for at least 6 generations in a C57BL/6N genetic background before the start of experimental procedures.

### Histology

Immunofluorescence and tissue processing were performed as previously described (66). In short, embryonic (E11.5 and E19) and postnatal brains (P0 and P56) were fixed in 4% paraformaldehyde (PFA), made in phosphate saline buffer (PBS), for 4 hours at 4°C. After fixation, embryonic and postnatal brains were cryoprotected in 30% sucrose in PBS, embedded and frozen in Tissue-Tek OCT (Sakura Finetek). Frozen brains were sectioned at 20 μm using a cryostat (Leica Biosystems). Sections were washed in PBS and blocked in PBS containing 5% normal goat serum (Sigma-Aldrich) (v/v) and 0.1% Triton X-100 (v/v) (Sigma-Aldrich) at room temperature for 2 hours. Sections were subsequently incubated in primary antibodies at room temperature overnight, washed in PBS and further incubated in secondary antibodies prepared in blocking solution for 3 hours at room temperature. The following antibodies were used: rabbit anti-cFos (1:3000; Cell Signaling Technology, 2250S), guinea pig anti-cFos (1:2000; Synaptic Systems, 226308), goat anti-GFP (1:1000; Rockland, 600-101-215); rabbit anti-GFP (1:1000; MBL International, 598), rabbit anti-HA (1:50; Chromotek, 7C9), guinea pig anti-Lbx1 (1:10000, ref. 32), guinea pig anti-Lmx1b (1:20000; ref. 32), goat anti-Phox2b (1:2000; R&D Systems, AF4940), goat anti-RFP (1:2000; Rockland, 200-101-379) and rabbit anti-RFP (1:2000; Rockland, 600-401-379). Cy2, Cy3 and Cy5 conjugated donkey anti-rabbit, anti-guinea pig, anti-goat secondary antibodies were obtained from Jackson ImmunoResearch and used at a concentration of 1:500. Fluorescence signals were acquired with: (i) a Zeiss LSM 700 confocal microscope using the automatic tile scan modus (10% overlap between tiles) and assembled using ZEN2012, (ii) a Zeiss spinning disk confocal microscope using the automatic tile scan modus (10% overlap between tiles) and assembled using ZEN2012, (iii) a Nikon Widefield Ti2 microscope using the automatic tile scan modus (10% overlap between tiles) and assembled using Nikon elements viewer, and iv) an Olympus BX51 epifluorescence microscope. All photographs were acquired in a non-blind manner. Photomicrographs were processed for brightness and contrast corrections using Adobe Photoshop 2020.

### Cell quantifications

Cell quantifications were performed in a non-blind manner as previously described (25). Briefly, dB2 neuron and other cell quantifications were performed in non- consecutive 20-μm-thick sections encompassing the complete anterior-posterior brainstem axis. On average 80-85 sections were obtained per animal and cells in every second section were bilaterally quantified and defined as subtotal of cells. The estimation of total number of cells was obtained by multiplying the subtotal of quantified cells by 2 (25).

### Brain clearing, lightsheet microscopy and analysis

*dB2-Tomato* brains were cleared using the CUBIC protocol as previously described (67). Briefly, brains were dissected and fixed overnight at 4°C in 4% paraformaldehyde made in PBS. After washing overnight in PBS, lipids were removed using Reagent-1 (25% urea, 25% Quadrol, 15% Triton X-100, 35% dH2O) at 37°C until brains were transparent (4 days). Brains were then washed overnight at 4°C in PBS to remove Reagent-1 and then placed into Reagent-2 (25% urea, 50% sucrose, 10% triethanolamine, 15% dH2O) at 37°C for refractive index matching (3 days). All reagents were acquired from Sigma-Aldrich. Once the brains were cleared, they were imaged using a Zeiss Lightsheet Z.1 microscope. 3D reconstructions, photos and videos were created with arivis Vision4D. Video editions were done using the commercial software Procreate® and LumaFusion (LumaTouch LLC).

### Unrestrained whole-body plethysmography for juvenile and adult mice

Breathing recordings for juvenile (P21) and adult (P56) mice in ambient air, hypercarbia or hypoxia were performed using previously reported protocols with minor modifications (42, 62, 68, 69). Mice were placed in Data Science International (DSI) whole-body plethysmograph chambers (601-1425-001) and habituated for at least one hour on the experimental day. Mice were then individually recorded (one per chamber). Each breathing recording included an initial 30-minute period of acclimatization (habituation) followed by a 20-minute period of respiratory recordings in ambient air to determine baseline respiration. Respiratory recordings were taken in thermostable conditions (32°C) as previously recommended (69). Breathing recordings were acquired with a FinePointe Whole-Body Plethysmograph Unit with gas switch capability (DSI; 271-0500-290). The unit provided the plethysmograph chambers with a constant 1 L/min airflow. Breathing waveforms were acquired with the New FinePointe Software (DSI; 271-0500-CFG). Body temperature and body weight were recorded for tidal volume estimation at the beginning of the respiratory recordings as previously described (42, 62, 69). Tidal volume estimations and respiratory cycle lengths were computed in the New FinePointe Software. Calculations for minute ventilation were obtained from the values of tidal volumes and respiratory cycle lengths using the New FinePointe Software. Protocols to induce hypercarbic and hypoxic responses in mice are published elsewhere (42, 62, 69), and schematically described in Figures 2A and 3C. In brief, following the 20-minute period to determine baseline respiration, mice were exposed to either a gas mixture of 21% O2, 8% CO2, balanced N2 (for hypercarbia) or 10% O2, balanced N2 (for hypoxia) for 10 minutes, followed by an additional 20-minute period of post-gas exposure. For determination of hypercarbic responses, the last five minutes of respiration in hypercarbia were compared to the last five minutes of respiratory recordings in ambient air (before gas exposure). For determination of hypoxic responses, the period encompassing 61-180 seconds (two minutes) from the start of the gas exposure were compared to the last two minutes of respiratory recordings in ambient air (before gas exposure).

### Unrestrained whole-body plethysmography for neonatal mice

Breathing recordings for neonatal (P7) mice in ambient air, hypercarbia or hypoxia were performed using the above-described protocols with the following modifications: individual mice were placed in whole-body plethysmograph chambers suitable for pups (DSI, 601-1426-001). The plethysmographic chambers included a thermo-controlled warm bed set at 37°C. The FinePointe Whole-Body Plethysmograph Unit was adjusted to provide a constant 0.5 L/min airflow.

### Head-out plethysmography for newborn mice

Breeding females were monitored daily for vaginal plugs. The day of vaginal plug identification was defined as E0.5. Pregnant dams were visually monitored for natural delivery from day E19 in intervals of 20 minutes. The onset of labor was recorded. On average pregnant dams completed labor within 20 minutes. Newborns were recorded withing 30 minutes from the onset of labor. The FinePointe Whole-Body Plethysmograph Unit was adjusted to provide a constant 0.5 L/min airflow and coupled to the DSI Digital Preamplifier (601-2401-001). The whole-body plethysmograph chambers suitable for pups were couple to the DSI head-out conversion kit (601-1533-001) and sealed with DSI latex collars (601-1533-002). The protocol used to induce hypercarbia in newborn mice is illustrated in Figure 6E. Every recording included a 10- minute acclimatization period followed by a 10-minute period to determine baseline respiration, mice were then exposed to 21% O2, 8% CO2, balanced N2 (for hypercarbia) for 5 minutes, followed by an additional 10-minute period of post-gas exposure. For determination of hypercarbic responses, the last three minutes of respiration in hypercarbia were compared to the last five minutes of respiratory recordings in ambient air (before gas exposure).

### Exclusion criteria

Plethysmograph chambers were covered with translucent red plastic covers. The experimenter visually monitored the recorded mice for potential movements during the analyzed periods. Movements were manually recorded, identified from the waveforms and excluded from further analysis. On average, movements represented <10% of the recording time sessions. Mice actively vocalize in the neonatal period (P0-P9). In previous work we defined vocal breathing by concurrent plethysmography and auditory recordings using an UltraSoundGate condenser microphone capsule CM16 (sensitive to frequencies from 20 Hz to 180 kHz) and Avisoft Recorder software (sampling rate, 250 kHz; format, 16 bit) from Avisoft Bioacoustics (70). Vocal breathing was excluded from the respiratory analysis. It represented <5% of the recording time sessions in newborn and neonates.

### Intraperitoneal CNO injections

CNO (HelloBio, HB6149) was dissolved in sterile saline at 10 mg/ml concentration. Mice received a single intraperitoneal injection of CNO at a concentration of 10 mg/kg and subsequently placed in the plethysmographic chambers for 10 minutes before the start of acclimatization periods and downstream respiratory recordings.

### Statistics

Statistical analyses were performed using Prism 8 (GraphPad). Data are plotted in scatter dot plots, violin plots or column dot plots with means and standard deviations.

Normal distributions of the data were tested using D’Agostino & Pearson, Anderson-Darling, Shapiro–Wilk and Kolmogorov-Smirnov tests. The statistical significance between group means was tested by one-way ANOVA, followed by Tukey’s post hoc test (for multiple-comparison tests), or two-tailed t-test (for pair comparison tests). Ridgeline plots were were done in R using RStudio (version 2022.12.0+353) and the ggridges package. Poincaré plots were acquired using Prism 8. Poincaré plot analysis, including SD1 and SD2 quantification, was performed with Python (v3.10.5) using the Pandas, Numpy and Matplotlib library via JupyterLab (v3.2.1). R and Python scripts can be obtained directly from. No statistical method was used to pre-determine the sample size. No randomization or blinding was used for *in vivo* studies.

## Supporting information

Supplementary_Figures_LRHM

## Author Contributions

KC and YX performed most of the research and equally contributed to this work. KC, YX and LRH-M designed research studies. KC, YX, AP, LA, MC and LRH-M carried out all respiratory recordings. KC, YX and EDL performed brain clearing, brain imaging and 3D reconstructions. KC, YX, AP, AK and LRH-M analyzed data. KC, YX and EGI performed immunostaining. KC, YX and LRH-M imaged immunofluorescence. MS, FMR, J-FB and HGN contributed with new reagents and new analytical tools. LRH-M supervised and directed the project. LRH-M prepared all figures and wrote the manuscript with the input from all co-authors.

## Acknowledgments

We thank Prof. Carmen Birchmeier (Max-Delbruck Center for Molecular Medicine) for providing the *Lbx1^Cre^* mouse line and antibodies against Lmx1b. We also thank Dr. Patricia Jensen (National Institute of Environmental Health Sciences) and Dr. Jean-François Brunet (Institut de Biologie de l’École Normale Supérieure) for sharing the *RC::FL-hM3Dq* and *Phox2b^FlpO^* mouse lines. YX is a recipient of a scholarship from the China Scholarship Council. The authors greatly thank Prof. Maria G. Castro (University of Michigan) and Prof. Alistair Garrat (Charite Universitätsmedizin Berlin) for a critical reading of our manuscript, and Dr. Pimrapat Gebert (Charite Universitätsmedizin Berlin) for support with statistical analysis. MS acknowledges support from “Fondation Pour l’Audition” (FPA RD-2017-5). This work was supported by the Fritz-Thyssen-Stiftung (grant # 10.20.1.004MN) and Deutsche Forschungsgemeinschaft (grant # 450241946), both granted to LRH-M.

## Notes

**Conflict of interest**. The authors declare that no conflict of interest exists for this study.

### Competing Interest Statement

The authors have declared no competing interest.

