## Supplementary_Figures_LRHM for "Genetic identification of novel medullary neurons underlying congenital central hypoventilation syndrome"

**Supplementary Figure Legends**

**Supplementary Figure 1. Analysis of breathing activity of *Control* and *dB2-Silence* mice.** (**A**) Quantification of minute ventilation, tidal volume, respiratory cycle length ($T_{TOT}$), and respiratory frequency, in *Control* and *dB2-Silence* mice before CNO treatment. Breathing recordings were taken in ambient air. The precise number (n) of analyzed mice is displayed in the brackets underneath the studied stages (P7, P21 and P56). (**B**) Quantification of standard deviation (SD) 1 and SD2 in *Control* and *dB2-Silence* mice at the indicated stages and conditions. Every dot represents the mean of individual animals. Significance was determined using one-way ANOVA followed by post hoc Tukey's analysis.

**Supplementary Figure 2. Hypercarbic response of *control* and *dB2-Silence* mice.** (**A**) Quantification of minute ventilation in *Control* and *dB2-Silence* mice before CNO treatment in ambient air (air) or hypercarbia ($CO_2$) as indicated in Figure 2A (upper panel). The precise number (n) of analyzed mice is displayed in brackets. Each dot represents the mean ± SD of the analyzed groups. (**B**) Respiratory responses to hypercapnia expressed as percentage of change relative to the baseline (ambient air). Change (Δ) of tidal volume observed in CNO-treated *Control* and *dB2-Silence* mice at the indicated ages. Please note this graph is related to Figure 2D. Every dot represents the mean of individual animals. Two-tailed t-tests was performed to determine statistical significance in A, and one-way ANOVA followed by post hoc Tukey's analysis in B.

**Supplementary Figure 3. Activation of dB2 neurons increases ventilation.** (**A**) Quantification of minute ventilation of *Control* mice before (gray) and after (dark blue) CNO treatment, while breathing ambient air at the indicated stages. (**B**) Respiratory changes in minute ventilation (left), tidal volume (middle), and respiratory cycle lengths ($T_{TOT}$, right) of CNO-treated *Control* and *dB2-Activity* mice while breathing ambient air. Changes (Δ) of the analyzed breathing parameters are expressed as percentage relative to the baseline (ambient air, before CNO treatment). (**C**) Representative plethysmographic traces of *dB2-Activity* mice before and after CNO at the indicated conditions and stages. (**D**) Quantification of minute ventilation of *dB2-Activity* mice in response to hypercarbia ($CO_2$, highlighted) before and after CNO treatment at the indicated stages. Respiratory recordings were taken in ambient air (air) or hypercarbia ($CO_2$) as illustrated in Figure 3C. Every dot represents the mean of individual animals. Significance was determined using one-way ANOVA followed by post hoc Tukey's analysis.

**Supplementary Figure 4. Lineage-tracing of neurons with a history of Lbx1 expression.** (**A**) Left, sagittal section taken from a *Lbx1*$^{Cre/+}$;*Rosa*$^{LSL\text{-}nGFP/+}$ mouse at birth (P0). The section was stained with GFP antibodies and DAPI. Right, same section as on the left, but displaying GFP signals in blue to allow for a better visualization of Lbx1 (red) immunoreactive neurons (magenta). Please note that this photomicrograph is also displayed in Figure 4C. (**B**) Left, transverse section taken from a *Lbx1*$^{Cre/+}$;*Rosa*$^{LSL\text{-}nGFP/+}$ mouse at birth. The section was stained with GFP antibodies and DAPI. Right, same section as on the left, but displaying GFP signals in blue to allow for a better visualization of Lbx1 (red) immunoreactive neurons (magenta). Note that at birth, most cells with a history of Lbx1 (GFP+) have already downregulated the active expression of this factor.

**Supplementary Figure 5. Distribution of Lbx1+/Phox2b+ brainstem neurons.** (**A**) Left, a transverse brainstem section at the level of facial motor (nV) nucleus stained with GFP antibodies and DAPI at birth (P0). The boxed area is shown on the right with Lbx1, GFP and Phox2b merged signals (in the middle), and Lbx1 and Phox2b only signals (on the right) for a better visualization of the Lbx1 and Phox2b double positive (magenta) cells. The small boxed area is magnified at the bottom. (**B**) Left, a transverse brainstem section at the level of nucleus ambiguus (NA) stained with GFP antibodies and DAPI at birth. The boxed area is shown on the right with Lbx1, GFP and Phox2b merged signals (in the middle), and Lbx1 and Phox2b only signals (on the right) for a better visualization of the Lbx1 and Phox2b double positive (magenta) cells. The small boxed area is magnified at the bottom. (**C**) Pie charts illustrating the proportion of cells in the intertrigeminal (ITR) and the peri nucleus ambiguus (periNA) groups with a history of Lbx1 expression (GFP+) and active expression of Lbx1 and Phox2b (dB2 neurons). (**D**) Quantification of Lbx1+/Phox2b+ (dB2) neurons in the intertrigeminal (ITR), vestibular (v1-v4), retrotrapezoid (RTN), epifacial (epiVII), and peri nucleus ambiguus (periNA), groups. Every dot represents the mean of individual animals. (**E**) Schematic transverse views of the mouse brainstem illustrating the location of Lbx1+/Phox2b+ (dB2) groups (magenta) at birth: i) intertrigeminal (ITR), vestibular (Ves), epifacial (epiVII), retrotrapezoid nucleus (RTN), and peri nucleus ambiguus (periNA) neurons. The cerebellum (cb), principal trigeminal nucleus (pV), trigeminal motor nucleus (nV), dorsal cochlear nucleus (DCN), nucleus tractus solitarius (nTS), spinal trigeminal nucleus (SpV), as well as the facial (nVII) and ambiguus (NA) motor nuclei are illustrated for anatomical orientation.

**Supplementary Figure 6. Intersectional labeling of dB2 neurons.** (**A**) Schematic view of a sagittal brainstem section illustrating the location of all dB2 neurons (in magenta) identified by intersectional genetics in this study. The genetic strategy is illustrated in Figure 5A. The area postrema (AP), nucleus tractus solitarius (nTS) as well as the facial (nVII) and ambiguus (NA) motor nuclei are illustrated for anatomical orientation. The blue lines indicate the transverse planes illustrated in B-D. Caudal dB2 groups (epiNTS, epiNA, periNA and infraSpV) are illustrated in Figures 5D and 5E. Analysis of ITR neurons was reported in ref. 12. (**C-D**) Main photographs, transverse brainstem sections stained with antibodies against the red fluorescent protein (to distinguish tdTomato) and DAPI. The boxed areas are magnified on the right of each main photograph illustrating tdTomato, Lbx1 and Phox2b merged signals. The small boxed areas are magnified at the bottom illustrating individual and merged fluorescent signals. The v2 marked group in C refers to the dB2 vestibular subgroup v2. Arrowheads in D point out to retrotrapezoid nucleus neurons. (**E**) Table illustrating the expression pattern of the identified dB2 neurons groups in this study.

**Supplementary Figure 7. Recombination patterns of *(Tg)Hoxb1$^{Cre}$*, *Egr2$^{Cre}$* and *(Tg)Hoxa3$^{Cre}$* driver lines.** (**A**) Sagittal brainstem sections taken from *(Tg)Hoxb1$^{Cre/+}$;Rosa$^{LSL\text{-}nGFP/+}$*, *Egr2$^{Cre/+}$;Rosa$^{LSL\text{-}nGFP/+}$* and *(Tg)Hoxa3$^{Cre/+}$;Rosa$^{LSL\text{-}nGFP/+}$* mice at birth (P0). Upper panels, the sections were stained with GFP antibodies and DAPI. The rhombomeres 3 to 6 are indicated. Lower panels, same sections as above displaying only GFP signals. The lines in *(Tg)Hoxb1$^{Cre/+}$;Rosa$^{LSL\text{-}nGFP/+}$* mice denote the transverse section planes illustrated in panels B-D. (**B-D**) Transverse brainstem sections taken from *(Tg)Hoxb1*$^{Cre/+}$*;Rosa*$^{LSL\text{-}nGFP/+}$, *Egr2*$^{Cre/+}$*;Rosa*$^{LSL\text{-}nGFP/+}$ and *(Tg)Hoxa3*$^{Cre/+}$*;Rosa*$^{LSL\text{-}nGFP/+}$ mice stained with GFP antibodies and DAPI at birth (P0).

**Supplementary Figure 8. Lineage-tracing of dB2 neurons using the *(Tg)Hoxb1*$^{Cre}$*, Egr2*$^{Cre}$ and *(Tg)Hoxa3*$^{Cre}$ driver lines.** (**A**) Transverse brainstem sections taken from *(Tg)Hoxb1*$^{Cre/+}$*;Rosa*$^{LSL\text{-}nGFP/+}$, *Egr2*$^{Cre/+}$*;Rosa*$^{LSL\text{-}nGFP/+}$ and *(Tg)Hoxa3*$^{Cre/+}$*;Rosa*$^{LSL\text{-}nGFP/+}$ mice stained with GFP, Lbx1 and Phox2b at birth (P0). Due to space restrictions the *Rosa*$^{LSL\text{-}nGFP/+}$ reporter is abbreviated only as *Rosa*$^{nGFP/+}$ in this figure. The boxed areas are magnified at the bottom of the main photographs. The facial motor nucleus (nVII), trigeminal motor nucleus (nV), principal trigeminal nucleus (pV) and nucleus ambiguus (NA) are indicated for anatomical orientation. (**B**) Quantification of the proportion of dB2 (Lbx1+/Phox2b+) neurons co-expressing GFP in each analyzed genotype, color code: *(Tg)Hoxb1*$^{Cre}$ (green), *Egr2*$^{Cre}$ (Maroon) and *(Tg)Hoxa3*$^{Cre}$ (blue).

**Supplementary Figure 9. Respiratory stability in *r4-Lbx1*$^{FS}$, *r3&5-Lbx1*$^{FS}$, and *r5&6-Lbx1*$^{FS}$ mice.** (**A**) Quantification of standard deviation (SD) 1 and SD2 in *Control*, *(Tg)Hoxb1*$^{Cre/+}$*;Lbx1*$^{FS/lox}$ (*r4-Lbx1*$^{FS}$), *Egr2*$^{Cre/+}$*;Lbx1*$^{FS/lox}$ (*r3&5-Lbx1*$^{FS}$), and *(Tg)Hoxa3*$^{Cre/+}$*;Lbx1*$^{FS/lox}$ (*r5&6-Lbx1*$^{FS}$) newborn (P0) mice. (**B**) Quantification of apnea lengths in of *Control*, *r4-Lbx1FS*, *r3&5-Lbx1FS*, and *r5&6-Lbx1FS* mice while breathing ambient air at the indicated stages. Each circle represents individual apneas. Significance was determined using one-way ANOVA followed by post hoc Tukey's analysis.

**Supplementary Figure 10. Loss of dB2 neurons in *r4-Lbx1*$^{FS}$*, r3&5-Lbx1*$^{FS}$*, and r5&6-Lbx1*$^{FS}$ mice.** (**A-C**) Histological analysis and quantification of vestibular (v1-v4), retrotrapezoid nucleus neurons (RTN) and epifacial (epiVII) dB2 neurons in *Control*, *(Tg)Hoxb1*$^{Cre/+}$*;Lbx1*$^{FS/lox}$ (*r4-Lbx1*$^{FS}$), *Egr2*$^{Cre/+}$*;Lbx1*$^{FS/lox}$ (*r3&5-Lbx1*$^{FS}$), and *(Tg)Hoxa3*$^{Cre/+}$*;Lbx1*$^{FS/lox}$ (*r5&6-Lbx1*$^{FS}$) newborn (P0) mice. Transverse brainstem sections were stained with Lbx1 and Phox2b antibodies, and counterstained with DAPI. The cochlear nucleus (Cn), spinal trigeminal (SpV), and facial motor nucleus (nVII) are indicated for anatomical orientation. Please note that the magnifications shown in Figure 6H were taken from the photographs illustrated here in panel A. (**D**) Schematic summary of the dB2 neuron distribution in the indicated genotypes. (**E**) Histological analysis and quantification of intertrigeminal region (ITR) and peri nucleus ambiguus (periNA) neurons in *Control*, *r4-Lbx1*$^{FS}$, *r3&5-Lbx1*$^{FS}$, and *r5&6-Lbx1*$^{FS}$ newborn (P0) mice. The transverse brainstem sections were stained with Lbx1 and Phox2b antibodies, and counterstained with DAPI. The trigeminal (nV) and ambiguus (NA) motor nuclei are indicated for anatomical orientation. Every dot represents the mean of individual animals. Significance was determined using one-way ANOVA followed by post hoc Tukey's analysis.

**Supplementary Figure 11. Respiratory dynamics of *r4-Lbx1*$^{FS}$*, r3&5-Lbx1*$^{FS}$*, and r5&6-Lbx1*$^{FS}$ mice in the postnatal life.** (**A**) Quantification of tidal volume, and respiratory cycle length ($T_{TOT}$) in *Control*, *r4-Lbx1*$^{FS}$, *r3&5-Lbx1*$^{FS}$, and *r5&6-Lbx1*$^{FS}$ newborns while breathing ambient air at P7 (top), P21 (middle) and P56 (bottom). Source data can be found in Supplementary Data 6. (**B**) Respiratory response to hypercarbia expressed as percentage of change relative to the baseline (ambient air).

Change (Δ) of minute ventilation, tidal volume and respiratory cycle length ($T_{TOT}$) displayed by the indicated genotypes at P7 (Top), P21 (middle) and P56 (bottom). Source data can be found in Supplementary Data 6. (**C**) Histological characterization of cFos activated cells in the midline raphe (raphe obscurus, pallidus and magnus), the nucleus tractus solitarius (nTS) and parabrachial complex (PBc) in *Control* and *r3&5-Lbx1$^{FS}$* mice, after an hour of hypercarbia exposure, at P56. The area postrema (AP) is also indicated for anatomical orientation. Note that the dorsal raphe (d-Raphe) also presented cFos activated cells after hypercarbic exposure. Quantification of cFos+ cells per each of the illustrated regions can be found in Figures 7D-7G.

**Supplementary Figure 12. Mis-specification of caudal dB2 neurons in the *Lbx1$^{FS}$* background.** (**A-B**) Left, sagittal schemas showing the transverse section planes illustrated on the right. Right, immunofluorescent characterization of caudal dB2 neuron groups (epiNTS, epiNA, periNA and infraSpV) in *dB2-Tomato* and *dB2-Tomato-Lbx1$^{FS}$* mice at embryonic (E) day 19. The transverse sections were stained against red fluorescent protein (to detect tdTomato), Lbx1 and Phox2b. The precise genotypes are display in brackets. The area postrema (AP), nucleus tractus solitarius (nTS), spinal trigeminal nucleus (SpV), as well as the vagal (nX), hypoglossal (nXII) and ambiguus (NA) motor nuclei are illustrated for anatomical orientation.

**Supplementary Figure 13. Characterization of the *(Tg)Hoxa3$^{Cre}$* and *MafB$^{Cre}$* expression patterns in early development.** (**A-B**) Characterization of the recombination patterns of the *Lbx1$^{Cre}$*, *Egr2$^{Cre}$*, *(Tg)Hoxa3$^{Cre}$* and *MafB$^{Cre}$* driver lines at E11.5, a time point in which dB2 neurons become specified. Recombination was assessed using the *Rosa$^{LSL\text{-}nGFP}$* reporter allele. This analysis included rhombomeres

5 to rhombomere 7. Please note that dA3 (Lbx1-/Phox2b+) neurons extend from rhombomere 4 to 7, while dB2 (Lbx1+/Phox2b+) neurons extend from rhombomere 2 to rhombomere 6, but not into rhombomere 7 (reviewed in refs. 31 and 35). (**A**) Transverse sections of the developing hindbrain for the indicated genotypes at E11.5. The sections were stained with antibodies against GFP, Lbx1 and Phox2b. The photographs are presented with Lbx1, GFP and Phox2b merged signals (on the left) and Lbx1 and Phox2b signals (on the right) for a better visualization of dA3 and dB2 neurons. (**B**) Schematic display of the recombination patterns observed in *Egr2*$^{Cre/+}$*;Rosa*$^{LSL\text{-}nGFP/+}$ (n=3), *(Tg)Hoxa3*$^{Cre}$*;Rosa*$^{LSL\text{-}nGFP/+}$ (n=4), and *MafB*$^{Cre}$*;Rosa*$^{LSL\text{-}nGFP/+}$ (n=3), embryos at E11.5. The blue dashed lines indicate consecutive sections (20 μm spaced between them) analyzed from the caudal border of rhombomere 6. In this analysis, we considered the caudal border of rhombomere 6 (denoted as 0 micrometers) to be the region in which Phox2b+ (dB2) progenitor cells were first seen (from the caudal to rostral axial level). The green bars indicate the recombination patterns seen in each of the analyzed embryos.

**Supplementary Figure 14. Respiratory patterns and histology of *MafB-Lbx1*$^{FS}$ newborn mice.** (**A**) Quantification of minute ventilation in *Control* and *MafB-Lbx1*$^{FS}$ newborns while breathing ambient air or hypercarbia. (**B**) Respiratory responses to hypercarbia expressed as percentage of change relative to the baseline (ambient air). Change (Δ) of minute ventilation (left), tidal volume (middle) and respiratory cycle length (right) displayed by *Control* and *MafB-Lbx1*$^{FS}$ newborns. (**C-D**) Histological characterization and quantification of vestibular (v1-v4) dB2 neurons as well as epifacial (epiVII) and retrotrapezoid nucleus (RTN) neurons in control and *MafB-Lbx1*$^{FS}$ newborns.

### Supplementary Figure 1

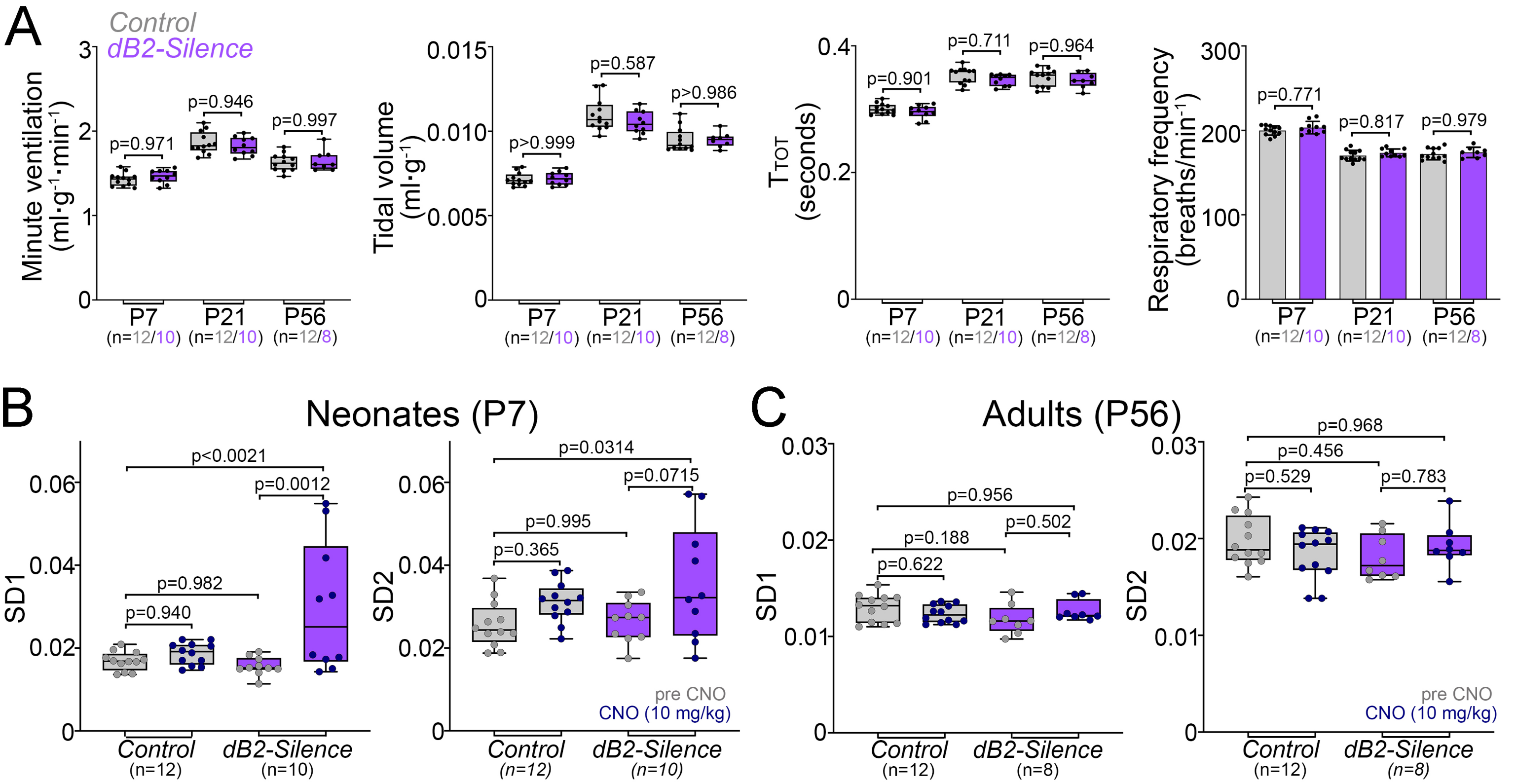

### Supplementary Figure 2

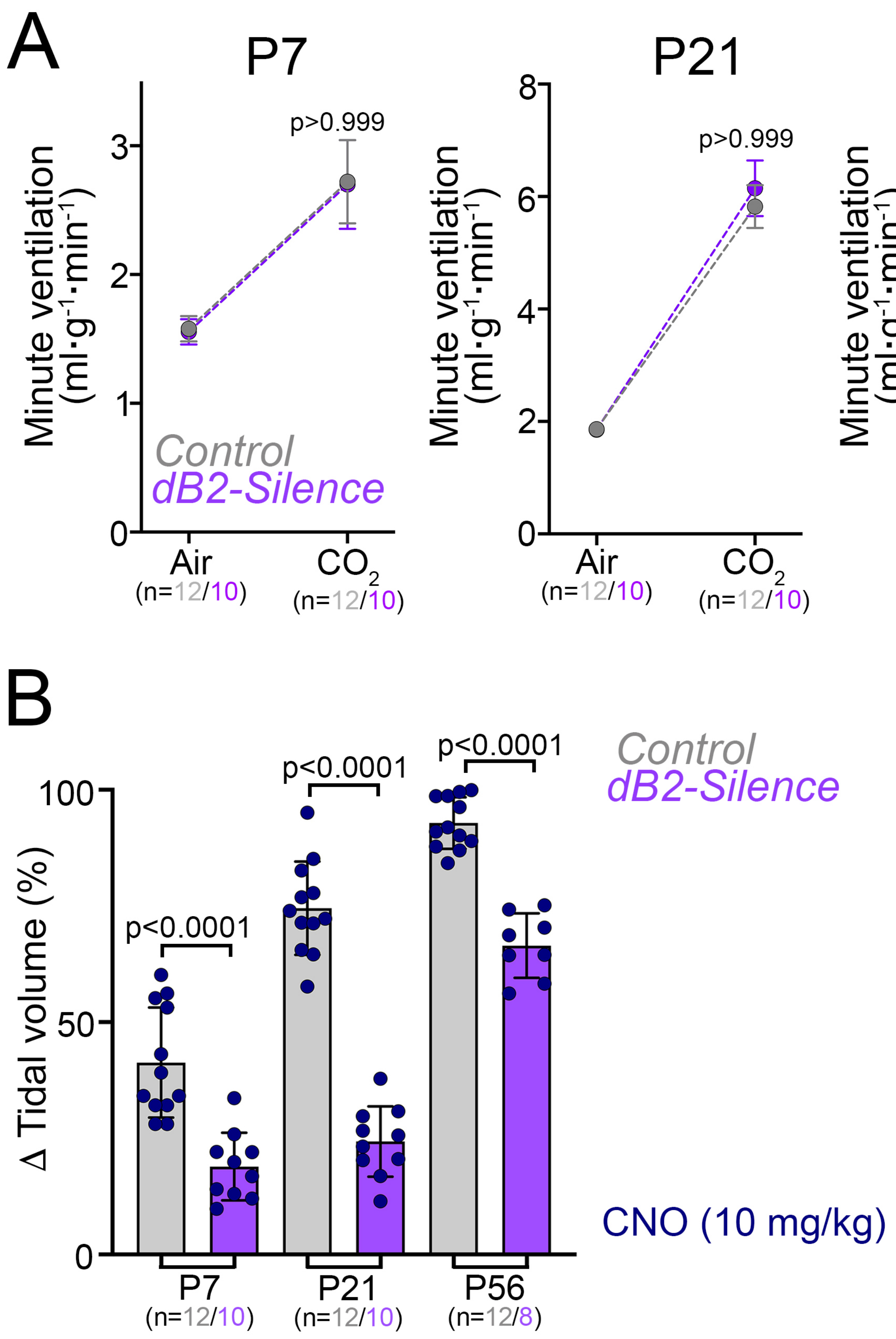

### Supplementary Figure 3

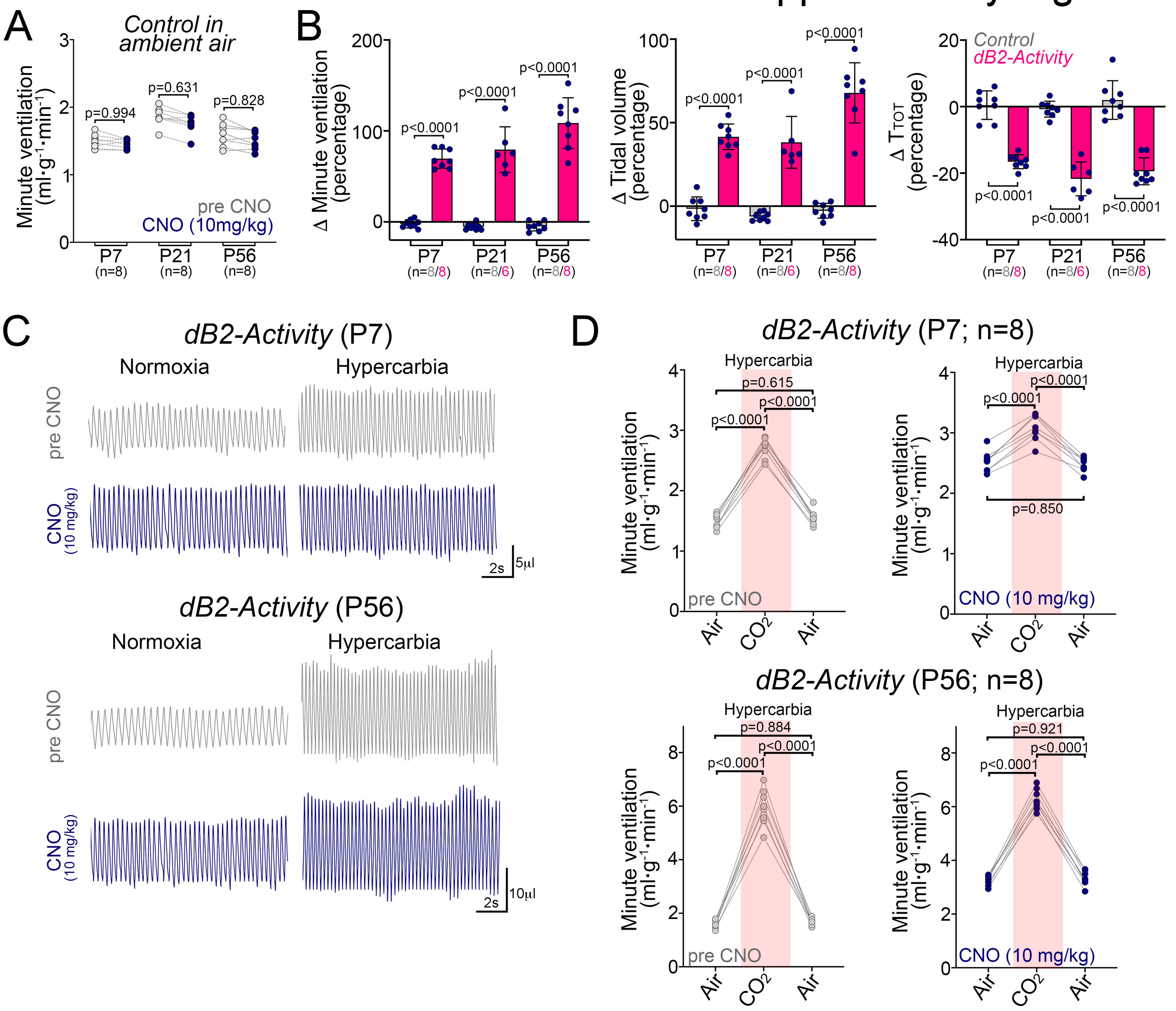

### Supplementary Figure 4

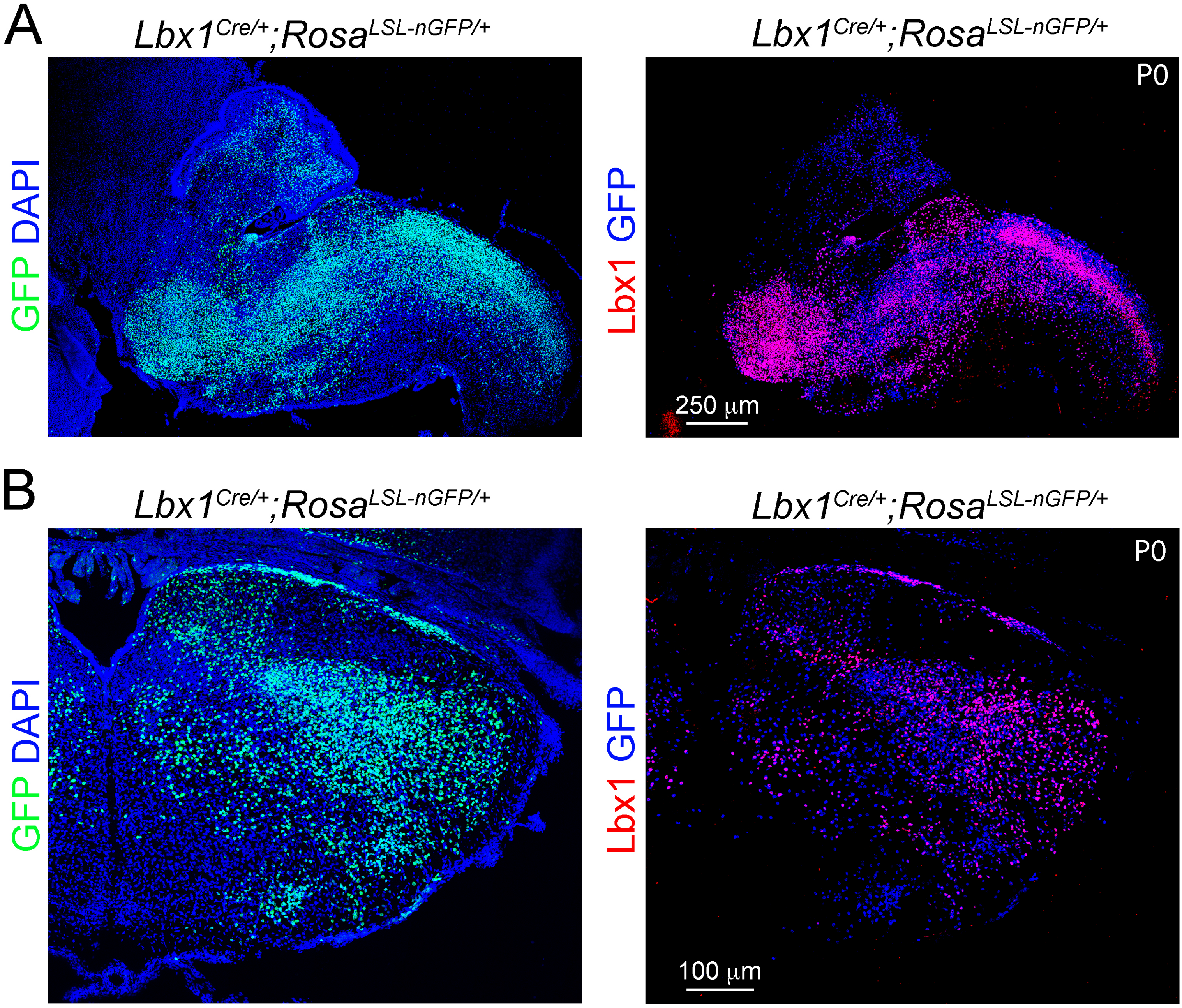

### Supplementary Figure 5

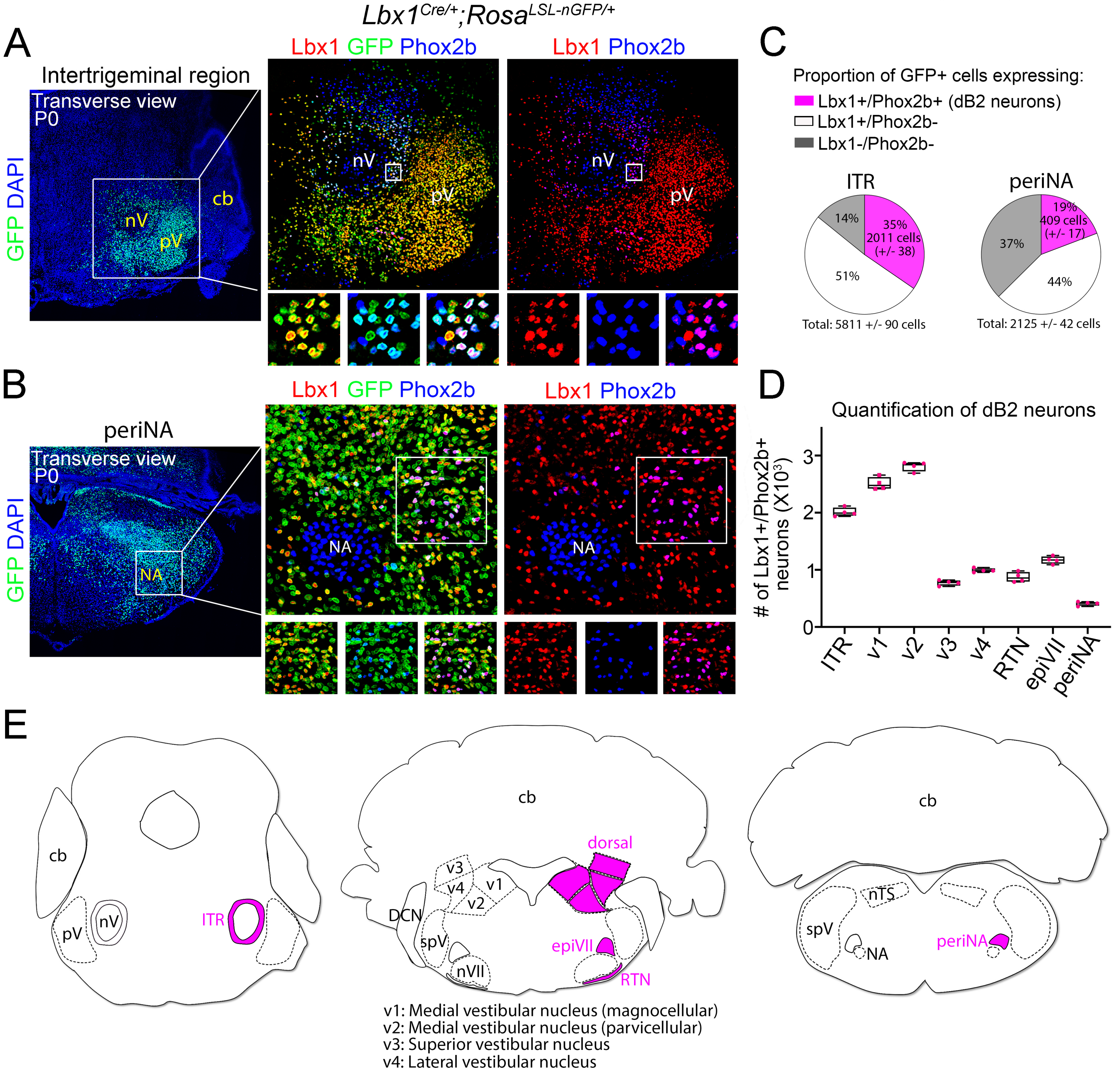

### Supplementary Figure 6

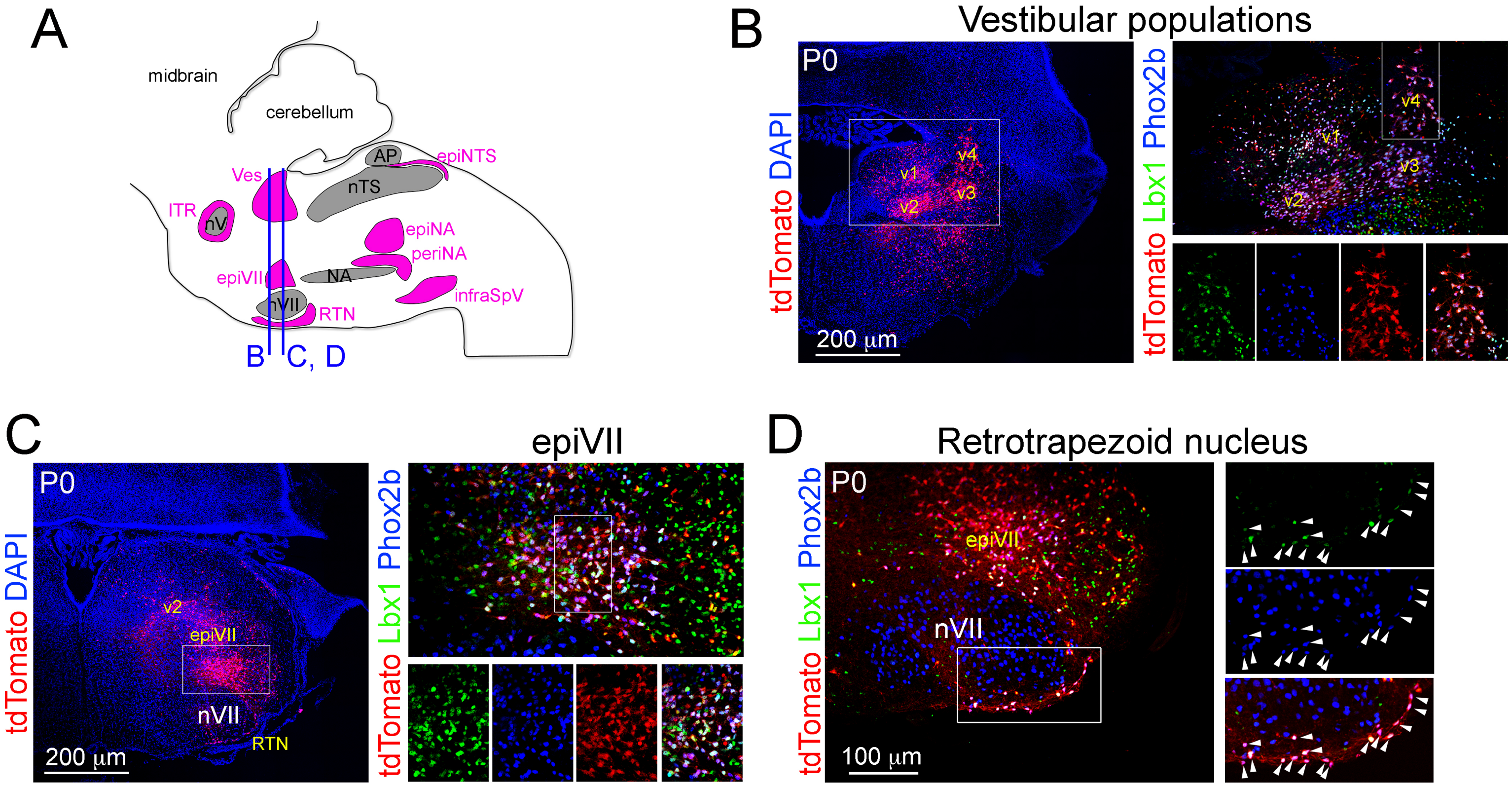


E

| dB2 neuron population | Active co-expression of Lbx1 and Phox2b |
|---|---|
| ITR | Yes |
| Ves (v1-v4) | Yes |
| epiVII | Yes |
| RTN | Yes |
| periNA | Yes |
| epiNA | No |
| epiNTS | No |
| infraSpV | No |

### Supplementary Figure 7

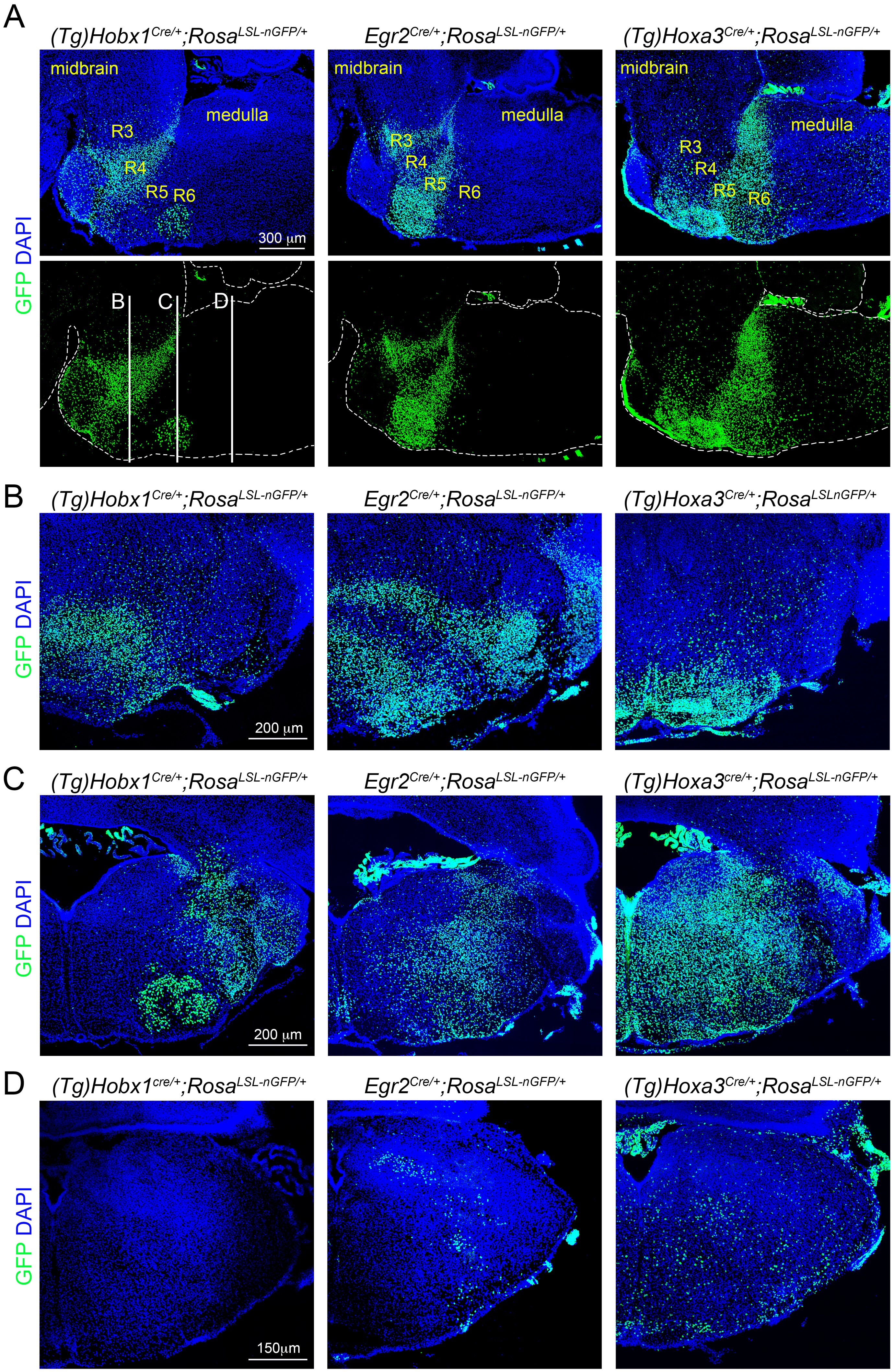

### Supplementary Figure 8

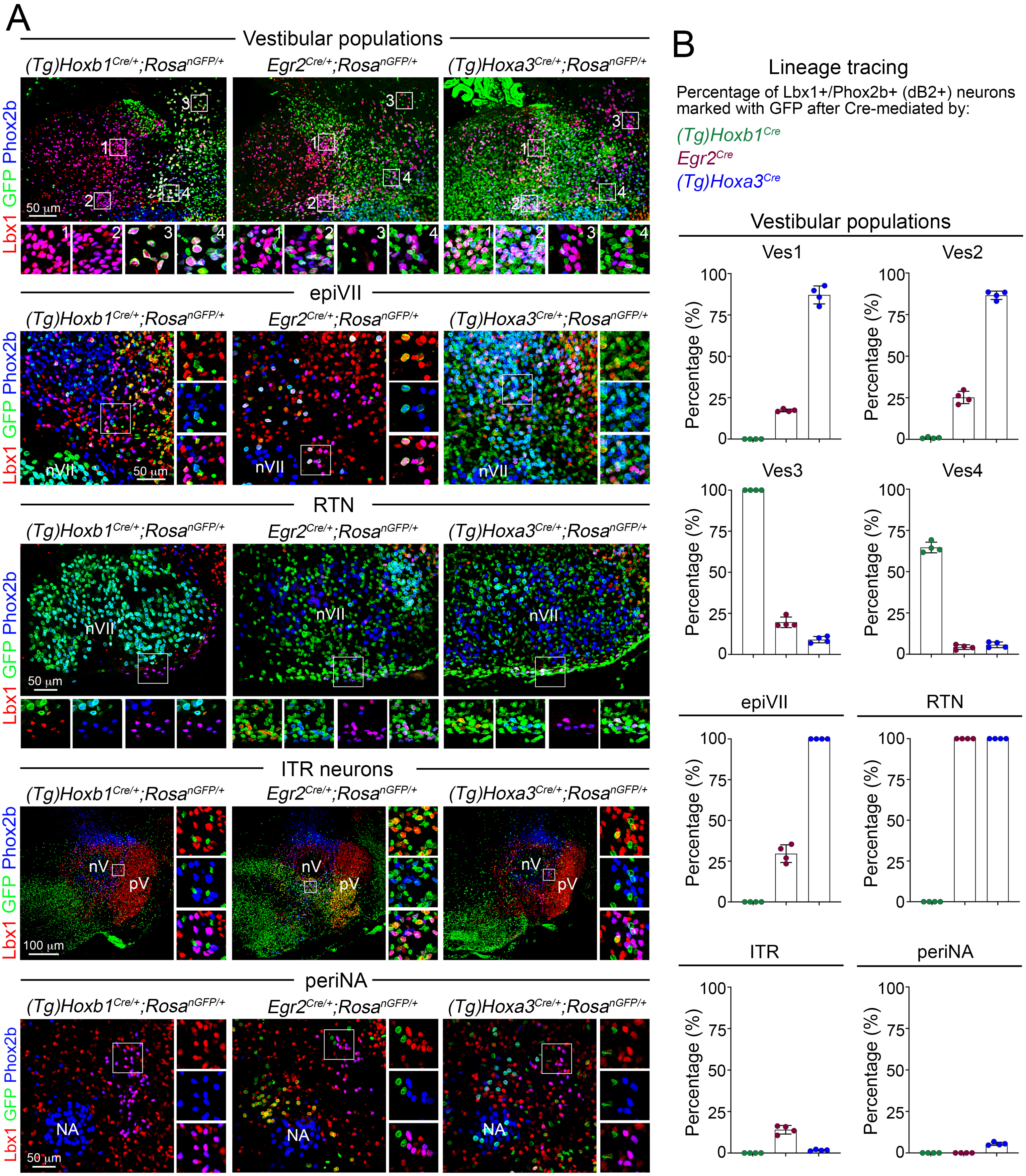

### Supplementary Figure 9

A

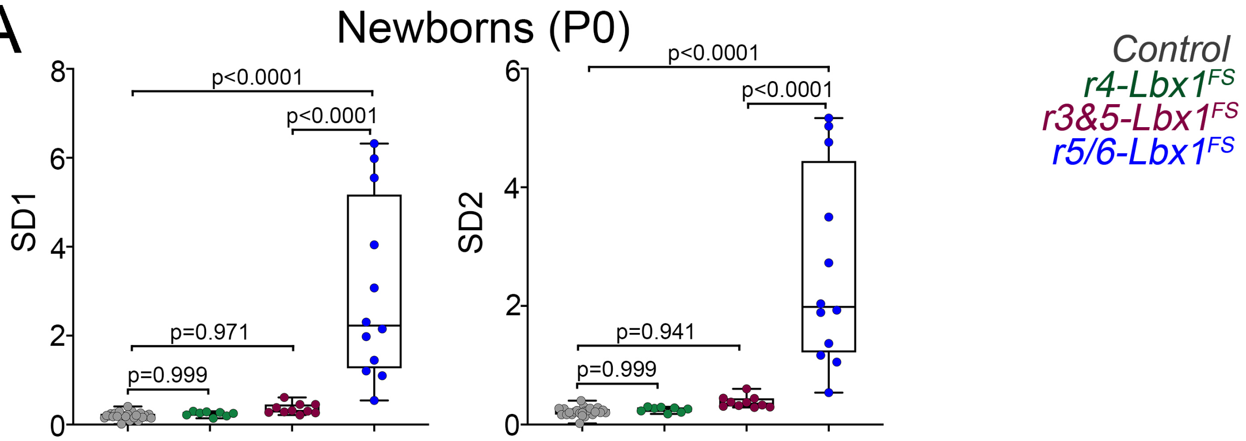


B

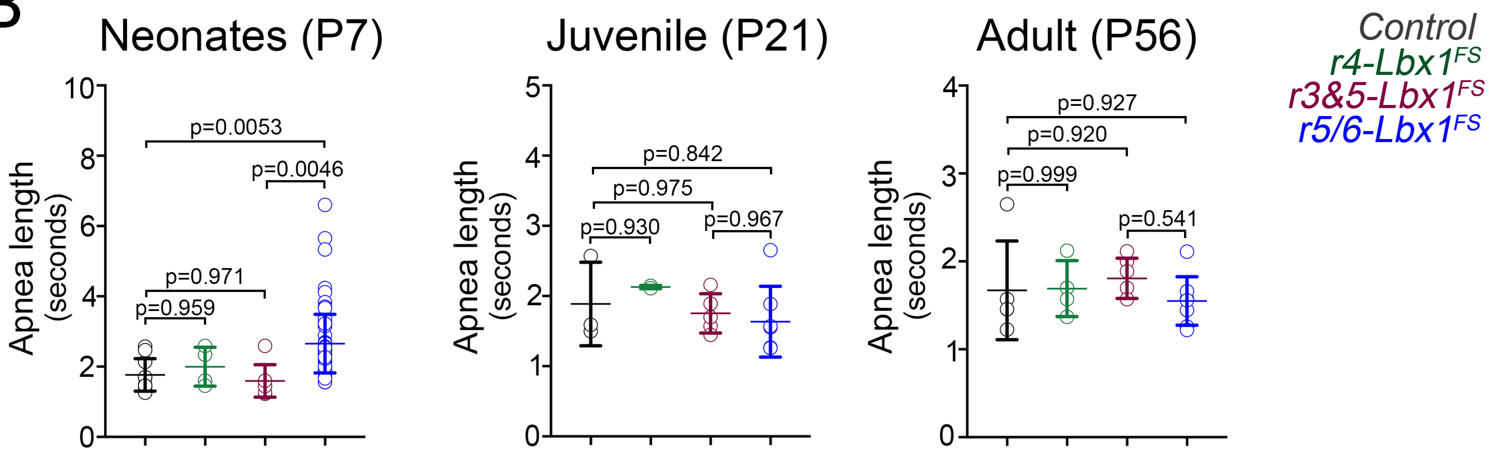

### Supplementary Figure 10

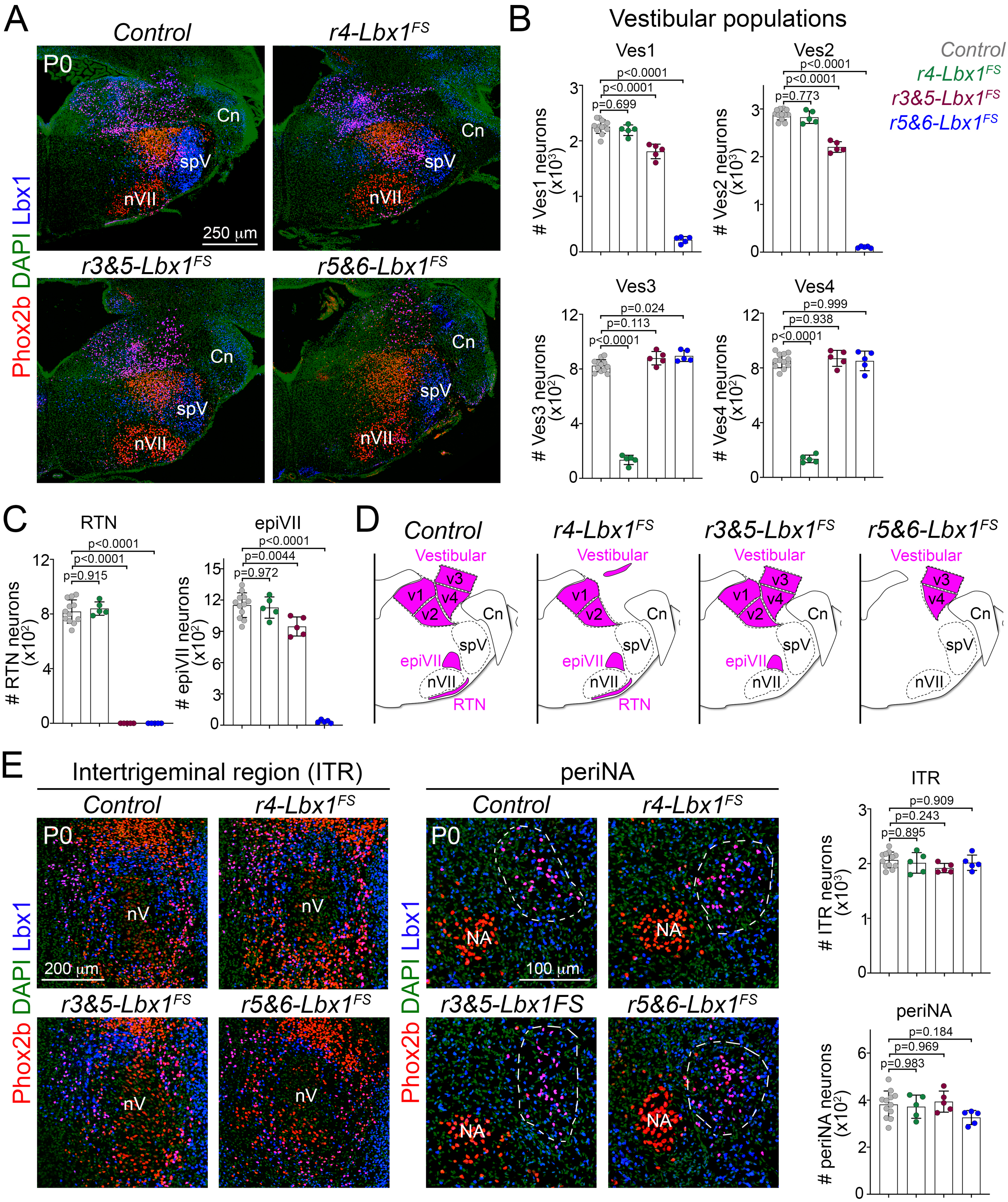

### Supplementary Figure 11

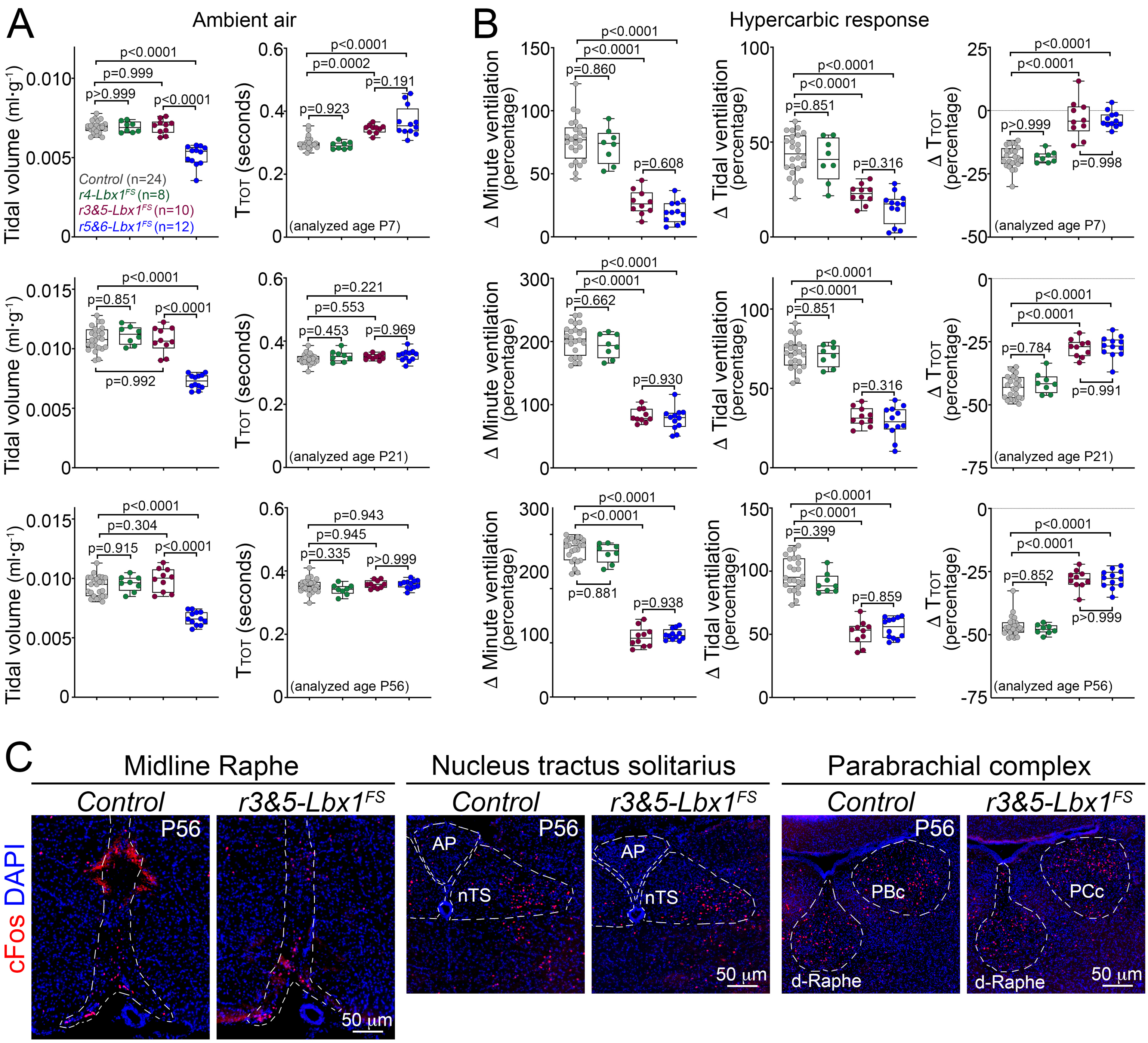

### Supplementary Figure 12

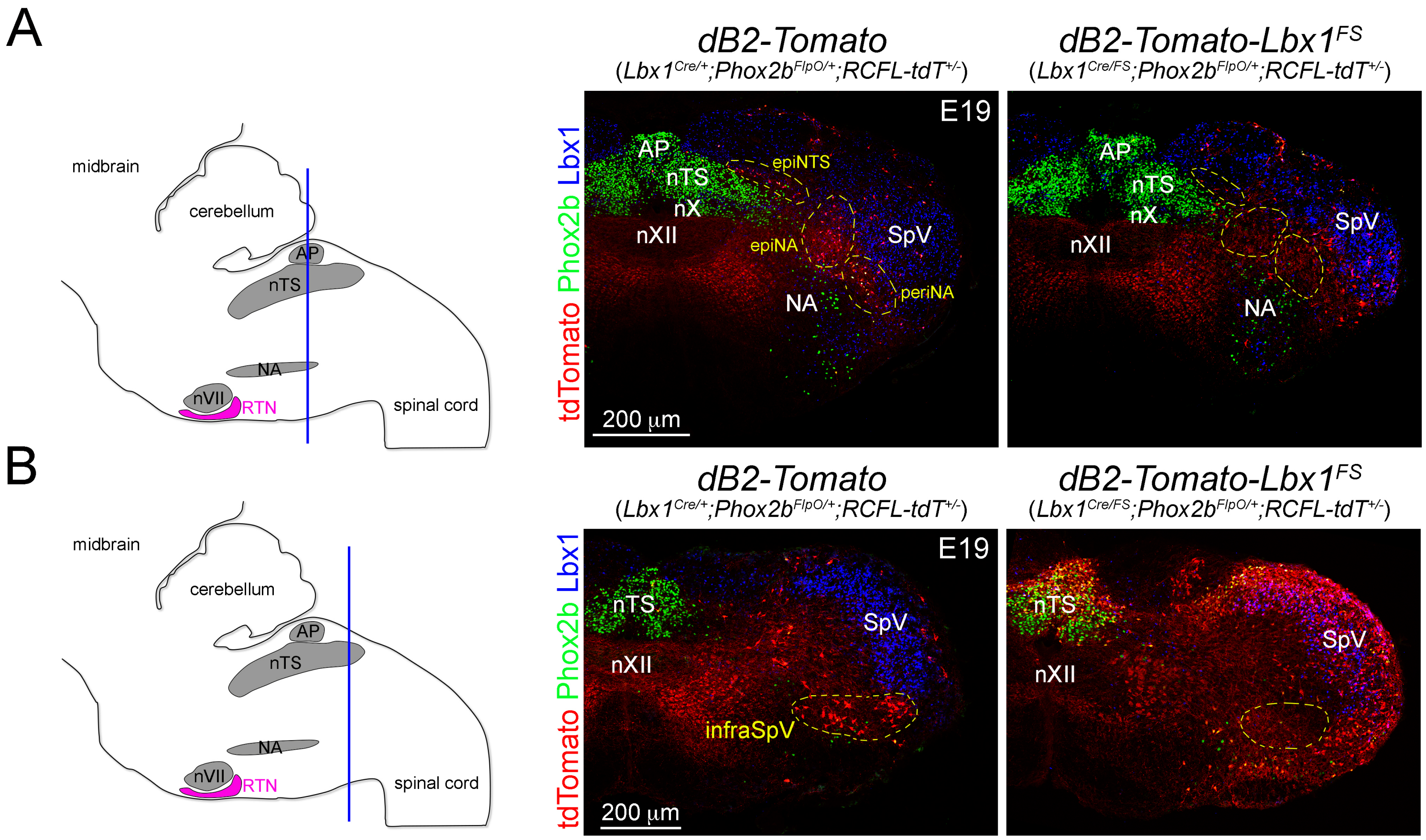

### Supplementary Figure 13

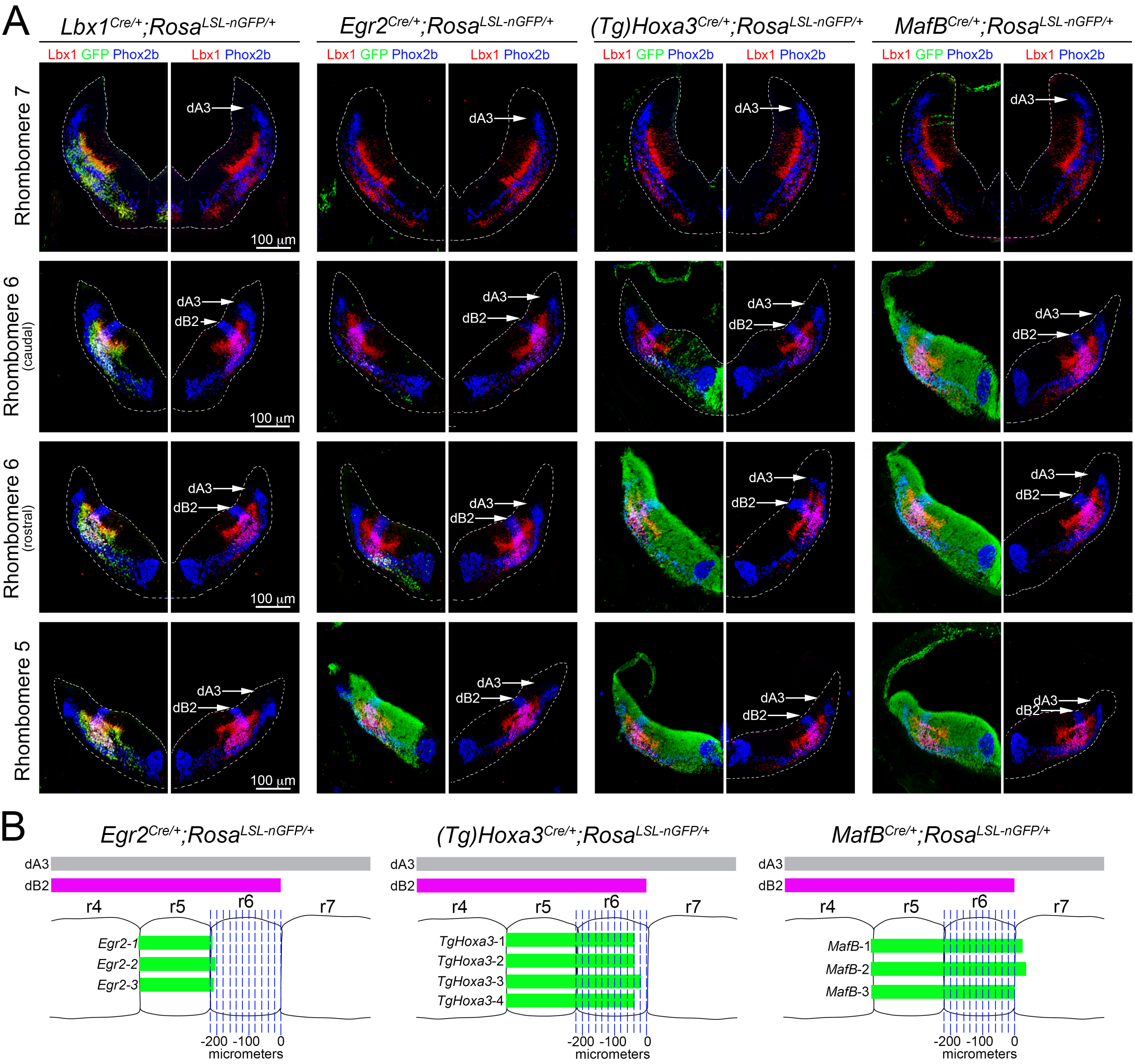

### Supplementary Figure 14

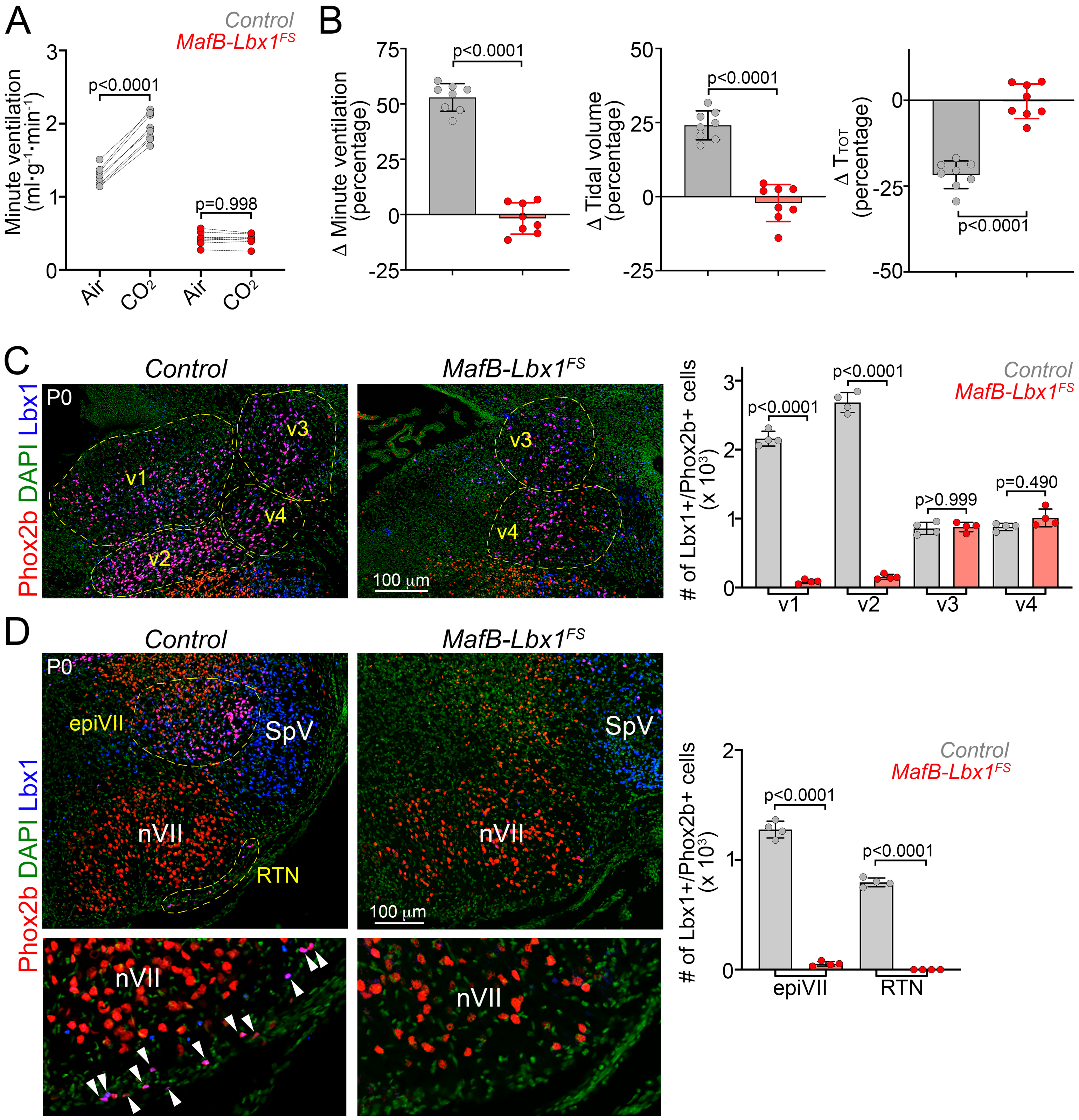
